# Dynamics of Cortical Dendritic Membrane Potential and Spikes in Freely Behaving Rats

**DOI:** 10.1101/096941

**Authors:** Jason J. Moore, Pascal M. Ravassard, David Ho, Lavanya Acharya, Ashley L. Kees, Cliff Vuong, Mayank R. Mehta

**Affiliations:** W.M. Keck Center for Neurophysics, Integrative Center for Learning and Memory, and Brain Research Institute; Neuroscience Interdepartmental Program, University of California at Los Angeles, U.S.A.; Department of Physics and Astronomy, University of California at Los Angeles, U.S.A.; Biomedical Engineering Interdepartmental Program, University of California at Los Angeles, U.S.A.; Departments of Neurology and Neurobiology, University of California at Los Angeles, U.S.A.

## Abstract

Neural activity *in vivo* is primarily measured using extracellular somatic spikes, which provide limited information about neural computation. Hence, it is necessary to record from neuronal dendrites, which generate dendritic action potentials (DAP) and profoundly influence neural computation and plasticity. We measured neocortical sub- and supra-threshold dendritic membrane potential (DMP) from putative distal-most dendrites using tetrodes in freely behaving rats over multiple days with a high degree of stability and sub-millisecond temporal resolution. DAP firing rates were several fold larger than somatic rates. DAP rates were modulated by subthreshold DMP fluctuations which were far larger than DAP amplitude, indicting hybrid, analog-digital coding in the dendrites. Parietal DAP and DMP exhibited egocentric spatial maps comparable to pyramidal neurons. These results have important implications for neural coding and plasticity.

**One Sentence Summary:** Measurement of cortical dendritic membrane potential for several days in freely behaving rats reveals disproportionate dendritic spiking and analog and digital coding.

---

Microelectrode techniques have enabled the long-term measurement of extracellular somatic action potentials in freely behaving subjects. However, somatic action potentials are brief (~1 ms), occur rarely (~1.5 Hz in principal neocortical neurons), and only represent the binary output of neurons, while the vast majority of excitatory synapses are localized on their dendritic arbors, each spanning more than 1000 μm (*1*). *In vitro* studies show that dendrites support local spike initiation of DAP (*2–13*) and back-propagating action potentials (bAP) initiated at cell somata (*5–8,10,14–17*). DAP endow neurons with greater computational power and information capacity (*9,18–22*) by turning dendritic branches into computational subunits with branch-specific plasticity (*11–13*).

Two-photon calcium imaging (*17, 23, 24*), sharp electrode (*2*), or patch-clamp techniques (*25*) can be used to study dendrites *in vivo*, but these suffer from critical drawbacks. Calcium imaging is not a direct measure of subthreshold membrane potential or dendritic sodium spikes, and lacks sub-millisecond resolution. Sharp electrode and patch-clamp techniques damage or rupture the membrane, thus limiting the recording duration (*26*) and altering *in vivo* neural dynamics including firing rates (*27*); they are ill-suited to record from thin dendrites. Additionally, these methods often require the subject to be anesthetized or immobilized, which alter neural dynamics and limit possible behavioral paradigms.

Extracellular recording techniques can also, at times, record intracellular-like signals. Sharp glass pipettes occasionally measure “quasi-intracellular” somatic recordings in anesthetized animals (*28–32*). These recordings can be stable for at most a few hours (*29*), and the recorded spikes have identical shapes as somatic action potentials (~1 ms width) (*29,31,32*). Quasi-intracellular recordings are proposed to arise from a region of high electrical resistance electrically isolating the membrane under the microelectrode, resulting in large amplitude, intracellular-like signals. Microelectrode arrays can achieve an “in-cell” recording configuration (*33,34*) when cultured neurons engulf the recording electrodes, yielding a large seal resistance and allowing measurement of the intracellular voltage. How might these phenomena be leveraged to investigate dendritic activity *in vivo*? We used chronically implanted tetrodes (*35,36*) (see Methods), which are often flexible and have a narrow profile to mitigate cell damage. Astrocytes and microglia can form a high impedance sheath around a tetrode and shield off the rest of the extracellular space (*37–39*). We hypothesized that this process can encapsulate a dendrite alongside the tetrode, such that the voltages at the tetrode tips approximate the intracellular (DMP), without penetrating the dendrite.

## Results

We implanted 9 rats with hyperdrives containing up to 22 individually adjustable tetrodes, as described previously (*35,36*) (see Methods). Tetrodes usually recorded standard extracellular signals, including thin (<1 ms) extracellular spikes of negative polarity with ~100 μV amplitude (Fig. 1a), presumably of somatic origin(*40*). However, in several instances (25 recordings made over 194 total tetrodes (13% success rate) in 21 out of 847 total recording days (2% of recording sessions) across 9 rats, see Methods) the signal was dramatically different, resembling an intracellular dendritic recording (Fig. 1b, Movie S1). These signals manifested overnight while all tetrodes were stationary. Here the signal amplitude was orders of magnitude larger, containing broad (>5 ms base width) positive polarity spikes with amplitudes of the order of thousands of microvolts (Fig. 1b, c). These signals could be discerned only when the data acquisition system settings with high dynamic range were used. These measurements were obtained at a median of 18 (between 6 and 55) days after surgery and discontinuation of any psychoactive drug, ruling out any possible effects of anesthesia.

**Fig. 1.**
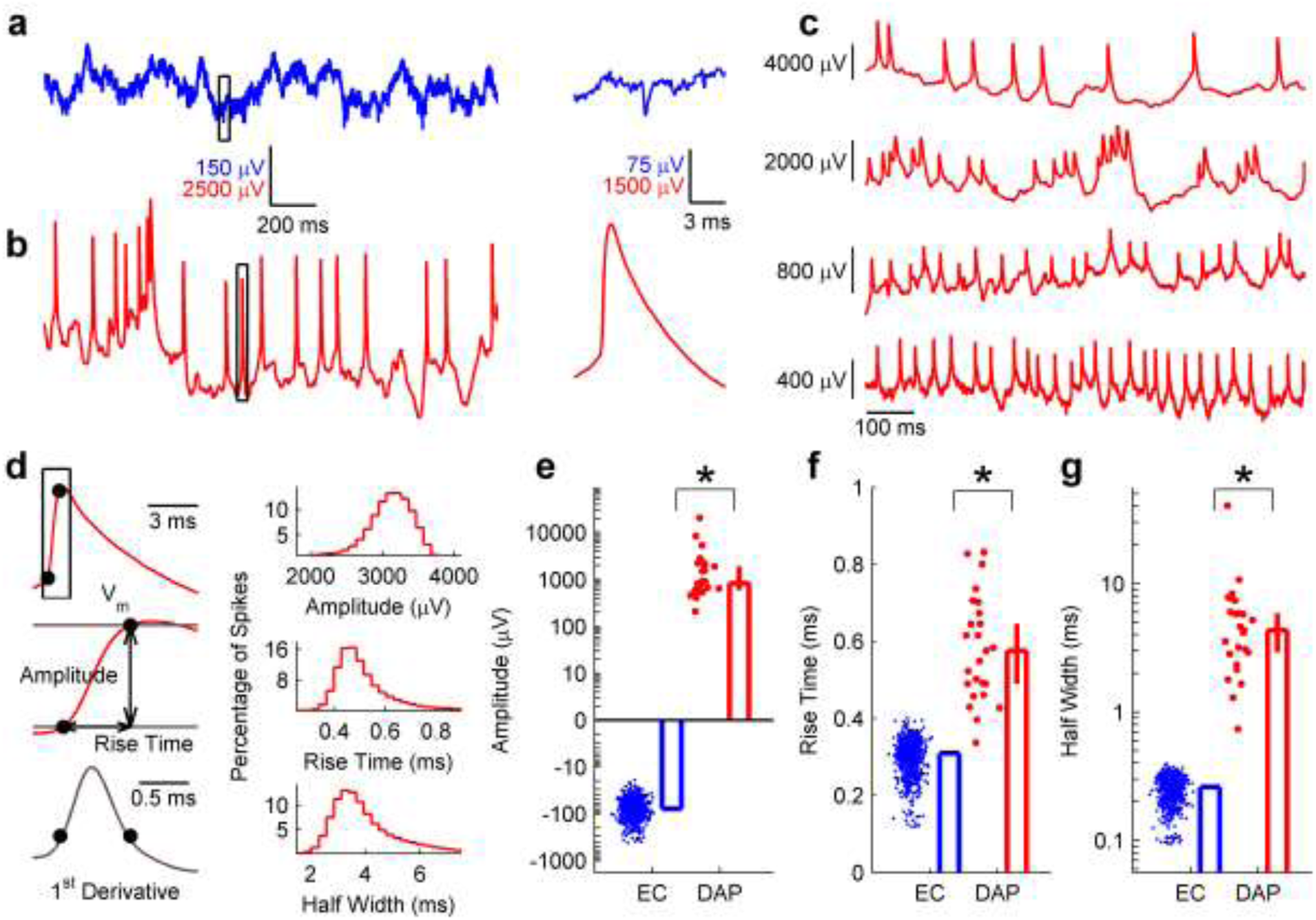
Measurements of DAP *in vivo*. **a**, Typical extracellular local field potential (LFP) showing ~100 μν fluctuations. Somatic action potentials are visible as thin, ~100 μV negative-polarity spikes (inset to right). **b**, Putative dendritic membrane potential recording on the same tetrode presented in a on the following day. Fluctuations are ~5000 μV. DAP are visible as broad, positive-polarity, ~5000 μV spikes with a much longer falling phase than rising phase (inset to right). See also Movie S1. **c**, Example membrane potential traces from four separate tetrodes, each exhibiting spontaneous dendritic spiking. **d**, Left, Quantification of DAP shape parameters (see Methods). Top right, distribution of DAP amplitudes within a single recording session, median 3120, [3110, 3130] μV, n=8187 spikes. Middle right, distribution of DAP rise times for the same recording session, median 0.49, [0.49, 0.49] ms, n=8187 spikes. Bottom right, distribution of DAP half widths for the same recording session, median 3.81, [3.78, 3.84] ms, n=8187 spikes. **e**, Extracellular spike amplitude (EC, blue) was always negative (−77.5, [−81.1, −73.4] μV, n=754 units), in comparison to DAP (red) which were always positive (835, [597, 190̅0] μV, n=25 dendrites), and more than 10 times larger in magnitude (p=1.7×10^−17^, Wilcoxon rank-sum test). **f**, The rise time of DAP (0.58, [0.49, 0.65] ms, n=25 dendrites) was significantly larger (p=1.7×10^−17^, Wilcoxon rank-sum test) than that of extracellular spikes (0.31, [0.31, 0.31] ms, n=754 units). **g**, The half-width of DAP (4.33, [2.92, 5.90] ms, n=25 dendrites) was also significantly larger (p=1.7×10^−17^, Wilcoxon rank-sum test) than that of extracellular spikes (0.26, [0.26, 0.26] ms, n=754 units). Data are reported and presented as median and 95% confidence interval of the median, and * indicates significance at the p<0.05 level.

## DAP vs extracellular spikes

For reasons described below, we refer to these spikes as DAP. We computed several features of the 25 DAP sources measured *in vivo* (Fig. 1d, see Methods), and compared these to extracellular somatic spike measurements from 754 units across 9 rats *in vivo* (Fig. 1e-g) and to available intracellular reports of dendritic spiking (table S1) (*3, 5, 7,8,10*). First, similar to in-cell and quasi-intracellular measurements (*29,30, 33, 34*), DAP amplitude was always positive (Median 850 μV, 25–75% range 570 to 2100 μV) (Fig. 1d, e, Fig. S1), in contrast to extracellularly recorded somatic spikes simultaneously recorded from nearby tetrodes; these were of negative polarity and of much smaller amplitude (Median −80 μV, range −40 to −400 μV) (Fig. 1e, Fig. S1). DAP rise time (0.57 ms,25–75% range 0.48 to 0.68 ms) was consistent across all recordings, similar to dendritic sodium spikes *in vitro*, and much larger than the rise time of the extracellular spikes (0.31 ms), but much faster than the rising phase of calcium spikes (Fig. 1d, f, table S1) (*3, 5,10, 23*). The full-width at half maximum (half-width) of DAP (4.3 ms, 25–75% range 2.7 to 6.3 ms) (Fig. 1d, g) was much longer than the DAP rise time (Fig. S2a, b), and far greater than the half-width of extracellular (0.26 ms, Fig. 1g) or intracellular (~0.7ms, table S1) somatic spikes; DAP were much wider than reported “high-amplitude positive spikes” from cortex(*41*) and somatic spikes recorded quasi-intracellularly (*28–32*). However, DAP half-width was much smaller than the typical half-width of dendritic calcium spikes (table S1). DAP width was more variable both within and across recordings (Fig. S2), in contrast to very consistent DAP rise times across data, and the two were significantly correlated (r=0.57, p=0.003) between DAP rise time and width across recordings (fig. S2f). These differences in the variability of rise time and width across DAP are also characteristic of dendritic sodium spikes (table S1) (*3, 5, 7, 8,10,14,17, 23, 24*).

## Mechanism of DMP recording, glial sheath hypothesis

We hypothesized that the DMP recordings were achieved by combining the above-mentioned in-cell (quasi-intracellular) recording (*29,30, 33, 34*) and glial-sheath mechanisms (*37,38*). Each tetrode bundle was ~35–40 μm in diameter, with a ~5 μm gap between tips. An intact dendrite, only a few microns thick (*1,10*), could thus be caught in this gap, and the tetrode would record an intracellular dendritic signal once the glial sheath formed. Immunohistochemistry of cortical sections previously implanted with tetrodes (see Methods) showed reactive astrocytes and microglia surrounding the tetrode location, with intact dendrites nearby (fig. S3a and Movie S2). Increased resistance from the electrode tips to ground due to this encapsulation would allow recording of the intracellular membrane potential (fig. S3b). We performed impedance spectroscopy measurements on electrodes during either DMP or local field potential (LFP) recordings (fig. S3c). Fitting parameters to an electric circuit equivalent model of glial encapsulation (*39,42*), the only parameter that was significantly higher (p=1.8×10^−3^) for DMP-recording electrodes was the resistance between the electrode tip and ground, through the glial sheath (fig. S3c, d). The estimated glial resistance was more than 6-fold larger for DMP-recording electrodes compared to LFP-recording electrodes; this is precisely the condition predicted by the model that would yield glial-sheath assisted measurements of the intracellular voltage (fig. S3b).

Further, we evaluated tetrode signal properties in the session before a DAP was recorded (PRE), during DAP recording (DUR), and in the session after the DAP signal was lost (POST). First, DAP amplitudes were nearly identical on all four channels of a tetrode in DUR (see Methods), unlike in the PRE condition where the voltages of extracellular spikes were different across four channels (Fig. 2a). This would not occur if any single electrode of the tetrode had penetrated the dendrite, but would be expected if the dendrite and the entire tetrode tip were encapsulated during DMP recording. Second, this would result in the shielding off of the surrounding extracellular medium from the tetrode by the encapsulating glial sheath. In the DUR condition both the amplitude and mean firing rate of detectable extracellular multi-unit activity (MUA) were significantly reduced (Fig. 2b, c, see Methods) compared to PRE. In one case, one of the four tetrode channels did not achieve an intracellular-like recording. MUA activity on only that channel was preserved, and its correlation with other channels remained low, suggesting that the single channel was not encapsulated by the glial sheath (Fig. S4). Third, there was no evidence of damage to the tetrodes that yielded DMP measurements, as all MUA properties in POST were similar to those in PRE (Fig. 2b, c).

**Fig. 2.**
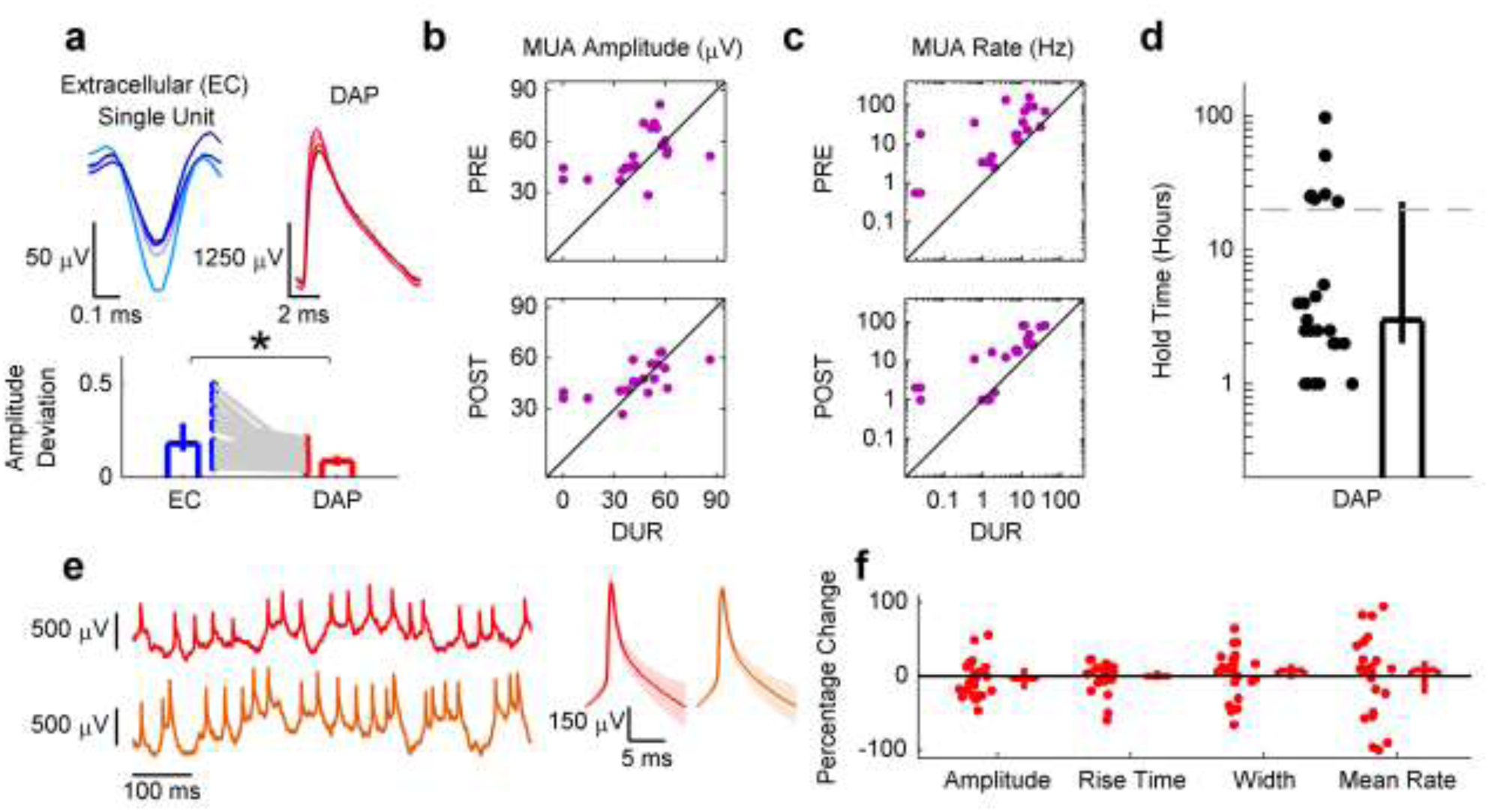
DAP measurements are similar across all electrodes of a tetrode and are stable for long periods. **a**, Top, the waveform of a single extracellular spike has different amplitudes on each of the 4 tetrode channels (left), but the waveform of a single DAP on the same tetrode the following day has very similar amplitudes (right). Bottom, the spike amplitude deviation across four electrodes (see Methods) was significantly greater for extracellular single units recorded the day before DAP were recorded (PRE) (0.18, [0.14, 0.29], n=25 units) compared to DAP during DMP recording (DUR) (0.09, [0.06, 0.12], n=25 DAP sources) on the same tetrode the following day (p=6.2×10^−6^, Wilcoxon signed-rank test) **b**, Multi-unit activity (MUA) amplitude during DMP recording (DUR) (41.8, [36.9, 53.4] μV, n=25 recordings) was significantly lower than during PRE (45.8, [44.3, 54.6] μV, n=25 recordings; p=6.6×10^−3^, Wilcoxon rank-sum test). MUA amplitude was smaller in DUR compared to the day after DMP recording (POST, 44.1, [40.6, 53.9] μV, n=25 recordings), but not significantly so (p=6.7×10^−2^, Wilcoxon rank-sum test). **c**, MUA rate was significantly lower in DUR (5.19, [1.29, 10.7], n=25 recordings) compared to both PRE (17.5, [4.84, 34.7] Hz, n=25 recordings; p=2.1×10^−5^, Wilcoxon rank-sum test) and POST (16.5, [2.01, 25.5] Hz, n=25 recordings; p=8.0×10^−5^, Wilcoxon rank-sum test). **d**, DAP recordings were stable for long periods of time (3, [2, 23] hours, n=25 recordings), with the shortest recording lasting 1 hour and the longest lasting 97.5 hours. Durations between 5.5 and 23 hours are absent due to restrictions on total recording duration in a single day, resulting in artificial bimodality. Hence, all recording durations are likely an underestimate of actual duration for which the DAP were held. e, Left, Membrane potential at the beginning of recording (top), and 90 minutes afterwards (bottom) of recording, showing little change in the quality of recording. Right, Averaged DAP (median and 25^th^, 75^th^ percentile) in the first part of the recording to the left (n=616 DAP) compared to the averaged DAP in the later part of the recording (n=897 DAP). f, Percentage change from the first five minutes of recording to the last two minutes was not significantly different from 0 for amplitude (−3.48, [−18.0, 11.7] % change, n=25 dendrites; p=0.55, Wilcoxon signed-rank test), rise time (0.00, [−5.00, 7.69] % change, n=25 dendrites; p=0.87, Wilcoxon signed-rank test), half-width (7.76, [−3.64, 16.27] % change, n=25 dendrites; p=0.35, Wilcoxon signed-rank test), nor mean firing rate (7.93, [−23.9, 20.5] % change, n=25 dendrites; p=0.97, Wilcoxon signed-rank test). Data are reported and presented as median and 95% confidence interval of the median unless otherwise noted, and * indicates significance at the p<0.05 level.

DAP properties remained stable for long periods of time (Fig. 2d-f). Our recordings typically lasted several hours, and DAP from the same tetrode could be recorded for up to 4 consecutive days (Fig. 2d, see Methods). From the beginning to the end of the recording span of any single DMP, there was no systematic change in DAP amplitude, rise time, width, or mean firing rate (Fig. 2e, f). These observations indicate that the tetrode did not damage the dendrite during these measurements (*26, 27,43,44*).

## Cellular identity of DAP

While the rise times and widths of the spikes described above are consistent with dendritically generated sodium spikes, i.e. DAP, they are also consistent with somatically generated spikes (bAP) (*5,7,8,10,14–17*). Hence, we compared the firing properties of DAP to those of extracellular somatic spikes recorded simultaneously on nearby tetrodes (see Methods). We first examined data from slow wave sleep (SWS), to minimize the influence of behavioral parameters (Fig. 3, see Methods). DAP mean firing rates were high (7.1 Hz), more than 4-fold greater than the mean firing rates of extracellular units (1.6 Hz) (Fig. 3a, b). The high DAP firing rates were not due to multiple DAP sources being pooled together, because all DAP had inter-spike interval and amplitude distributions inconsistent with multiple independent sources (Fig. S5, see Methods). The high rates were also unlikely to be due to damage or otherwise altered activity of dendrites because of the longevity of recording (Fig. 2d), and the absence of any systematic changes in DAP kinetics or rate (Fig. 2e, f) over long periods of time.

**Fig. 3.**
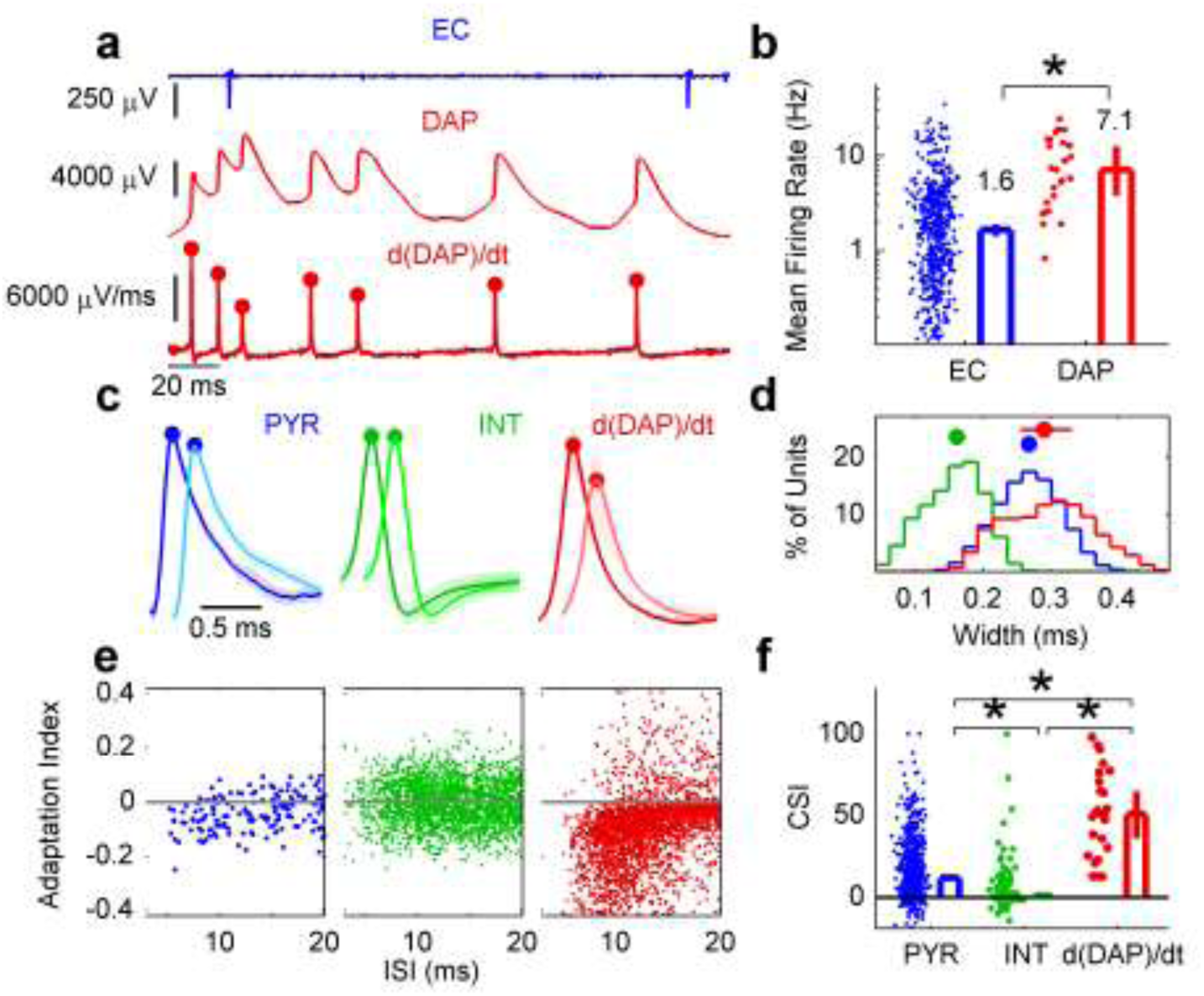
DAP are likely to be from pyramidal neurons but have much greater firing rates and stronger short-term plasticity. **a**, Top, sample LFP showing a single extracellular unit firing at a relatively low rate. Middle, sample putative membrane potential showing a DAP firing at a higher rate. Bottom, first temporal derivative of the MP trace. Red dots indicate the peak value of identified DAP. Note activity-dependent attenuation. **b**, In SWS, the mean spontaneous firing rate of extracellularly recorded units (1.65, [1.41, 1.91] Hz, n=754 units) was more than 4-fold smaller (p=4.1×10^−8^, Wilcoxon rank-sum test) than the firing rate of DAP (7.07, [3.76, 12.6] Hz, n=25 dendrites). Notice the near complete absence of low (<1 Hz) firing rate DAP. **c**, Demonstration of extracellular waveforms The median and 25^th^, 75^th^ percentile of sample pyramidal neuron (PYR, blue, n=140 spike pairs) interneuron (INT, green, n=6987 spike pairs) and d(DAP)/dt (red, 130 spike pairs) waveforms are plotted for the first spike (darker color) and the second spike in burst pairs (lighter color). PYR and DAP are broad, and the second spike has a smaller amplitude. This is not the case for INT. **d**, Extracellular spike width of PYR (0.27, [0.26, 0.27] ms, n=657 units) and DAP (0.29, [0.25, 0.33] ms, n=25 dendrites) were significantly greater than INT (0.16 [0.15, 0.17] ms, n=97 units; PYR vs INT, p=1.2×10^−52^; DAP vs INT, p=5.1×10^−14^, Wilcoxon rank-sum test for both). DAP were also significantly wider than PYR (p=1.5×10^−2^, Wilcoxon rank-sum test). **e**, Adaptation index (see Methods) plotted against inter-spike interval (ISI) for a sample pyramidal neuron, interneuron, and DAP. For PYR and DAP, but not INT, the adaptation index is largely negative for ISI less than 20 ms. For clarity, every 5^th^ spike pair is plotted for INT and DAP. **f**, PYR CSI (12.0, [10.3, 13.5], n=657 units) was significantly greater (p=1.0×10^−11^, Wilcoxon rank-sum test) than that of INT (1.02, [0.25, 2.99], n=97 units), but smaller (p=2.9×10^−10^, Wilcoxon rank-sum test) than that of DAP (50.8, [36.0, 65.0], n=25 dendrites). DAP CSI was significantly greater (p=4.7×10^−12^, Wilcoxon rank-sum test) than that of INT. Data are reported and presented as median and 95% confidence interval of the median unless otherwise noted, and * indicates significance at the p<0.05 level.

Could the high DAP rates be a result of preferentially recording from high-rate interneurons (*45*)? Interneurons have narrow spikes that exhibit very little spike amplitude attenuation within high frequency bursts, characterized by very low complex spike index (CSI) (*35,45–48*) (see Methods). In contrast, pyramidal neurons, have wide spikes that exhibit consistent amplitude attenuation. These distinctions between cell types are present in dendrites as well (*10,15,16, 49,50*). Thus, DAP should have wide spikes and a high CSI if recorded from pyramidal neurons.

We separated the extracellular units into putative pyramidal neurons and interneurons using standard methods (fig. S6a) (*45*) and compared their widths and CSI to the first temporal derivative of DAP, as an approximation of the extracellular waveform (*40,51*). Our data set contained far more pyramidal neurons (87%) than interneurons (13%). As expected, pyramidal neurons had significantly lower firing rates and larger widths and CSI than the interneurons (*45*) (Fig. 3c-f, fig. S6b) (*45,46, 52*). The estimated extracellular widths of DAP were slightly larger than pyramidal neurons but about twice as large as interneurons (Fig. 3d). The DAP CSI was slightly greater than pyramidal neurons’ but fifty-fold greater than interneurons (Fig 3e,f). The distributions of both of these DAP measures were unimodal and most similar to pyramidal neurons, indicating that our DAP were measured from the dendrites of pyramidal neurons.

DAP mean rate was significantly greater (more than five-fold) than that of pyramidal neurons (p=1.3×10^−9^) (fig. S6b), as was the peak firing rate (93.5 Hz DAP, 67.6 Hz pyramidal, p=1.0×10^−2^, fig. S6c). The differential pattern of the mean and peak rates might be explained by differences in short term plasticity (*46*) which is greater in the pyramidal neuron dendrites compared to soma(*15,16*). This is manifested in activity-dependent adaptation of DAP amplitude, rise time and width, similar to those in pyramidal neuron dendrites *in vitro* (*15,16*) (Fig. S6d, e).

To determine whether these are bAP or DAP and the locus of their measurement, we compared our experimental results with the simulations of a multicompartment model of cortical layer 5 pyramidal neuron (fig. S7)(*53–56*). The rise time and widths of our data had a significant overlap with those of simulated DAP generated in the distal-most dendrites and measured locally (fig. S7b, d). In contrast, these measured parameters had only a partial overlap with the bAP (fig. S7c, d). These results suggest that for the case of layer 5 pyramidal neurons, most of our measured DAP are locally generated and measured in the distal-most dendrites. Similar results may hold for other pyramidal neuron types and dendrites. This hypothesis is further supported by the observation that DAP rates are several fold greater than somatic rates (Fig. 3b, fig. S6b).

Further analysis of DAP properties did not show evidence of two different populations. The DAP rates were similar at all depths (from 330 μm to 1240μm, p=0.24, fig. S8a) and were greater than somatic rates at all depths (fig. S8b). Other properties of DAP sources were uncorrelated with DAP rate as well, including amplitude (p=0.19), rise time (p=0.14), half-width (p=0.49), and CSI (p=0.57) (fig. S8c), suggesting no clear demarcation between high and low rate DAP. The disparately large rates of DAP compared to somatic spikes suggests that only a fraction of DAP elicit somatic spiking, indicating a significant decoupling of dendrites and somata (*2, 6, 7,10,16*).

## Subthreshold dendritic potential dynamics during SWS

Subthreshold fluctuations accompanying DAP during SWS had characteristics suggesting they represent the DMP (Fig. 4a, see Methods). The subthreshold fluctuations were of the order of thousands of microvolts, reaching up to 20 mV, (Fig. 4a, b), far exceeding the range of the LFP on nearby tetrodes (Fig. 4b). The range of the subthreshold fluctuation was positively correlated with the DAP amplitude on the same recording (Fig. 4c). Nearby LFP range was not correlated with DAP amplitude (fig. S9a). These results could arise due to the quality of glial-sealed encapsulation (Fig. S3, 4). The subthreshold range was always greater (6.8-fold) than the corresponding DAP amplitude (Fig. 4c), a membrane property primarily observed *in vitro* in dendrites that are electrotonically distant from the soma (*3–8,10*). In all but one case, the DMP signal in SWS was significantly negatively correlated with the LFP signal on a nearby tetrode (Fig 4d, fig. S9b, Movie S1) similar to somatic membrane results (*57*).

**Fig. 4.**
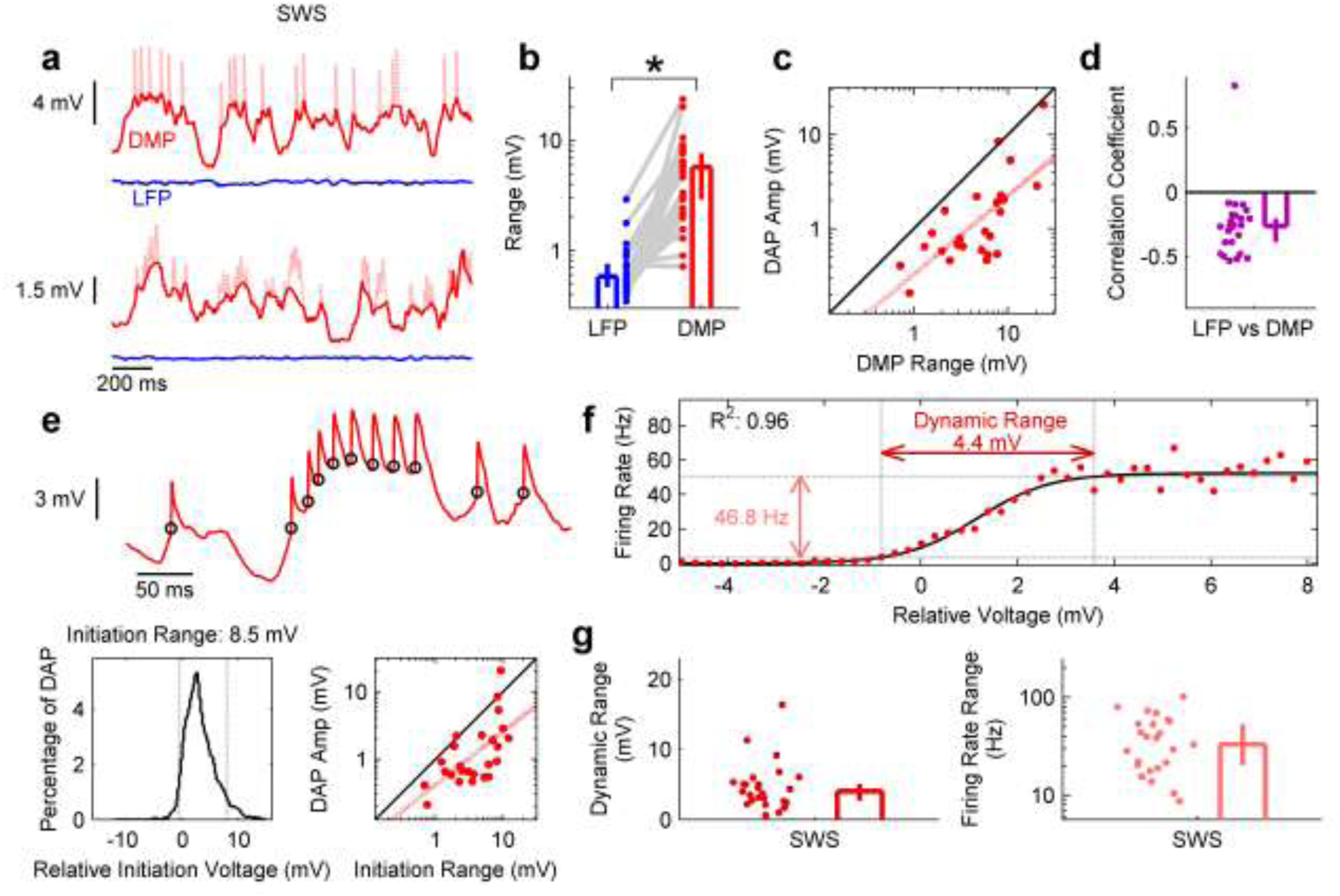
Large subthreshold membrane potential fluctuations modulate DAP rates during SWS. **a**, Top, sample dendritic membrane potential (DMP, red) trace during SWS, showing prominent oscillations of the same order of magnitude as DAP. DAP are shown in light red to highlight the spike-clipped subthreshold membrane potential. Below on the same scale is the local field potential (LFP, blue) recorded simultaneously on a nearby tetrode. Bottom, same as above but from a different pair of tetrodes in a different recording session. **b**, The range of the LFP (0.58, [0.47, 0.76] mV, n=25 LFP) was nearly an order of magnitude (9.9-fold) smaller (*: p=1.6×10^−8^, Wilcoxon signed-rank test) than the subthreshold DMP accompanying DAP (5.72, [2.92, 7.69] mV, n=25 dendrites). **c**, In SWS, the range of subthreshold DMP was nearly always larger (p=1.4×10^−5^, Wilcoxon signed-rank test) than the corresponding DAP amplitude (0.84, [0.60, 1.90] mV, n=25 dendrites), and positively correlated (r=0.71, [0.44, 0.86]; p=6.9×10^−5^, two-sided *t* test). **d**, In all but one case, LFP and simultaneously recorded DMP during SWS were negatively correlated (r=−0.26, [−0.38, −0.20]). See also Movie S1. **e**, Top, sample DMP trace segment showing a dynamic threshold for DAP initiation, with the initiation points marked by black circles. Bottom left, histogram of initiation voltages for entire recording session from which the above was taken; the 5–95% range of initiation voltages spans 8.5 mV. Bottom right, DAP initiation range (3.67, [2.24, 6.99] mV, n=25 dendrites) was larger (p=5.1×10^−5^, Wilcoxon signed-rank test) than the corresponding DAP amplitude, and positively correlated (r=0.63, [0.31, 0.82]; p=7.4×10^−4^, two-sided *t* test). **f**, Sample DAP firing rate as a function of relative voltage, which was well-approximated (fig. S9g) by a sigmoidal logistic function (black line). Firing rates here vary by 46.8 Hz over a dynamic range of 4.4 mV. **g**, Left, the population of DAP had a large dynamic range of initiation voltages (4.03, [2.74, 5.7] mV, n=25 dendrites), as defined in **f**. Right, the firing rate range of the DAP population was similarly wide (33.1, [20.3, 53.1] Hz, n=25 dendrites). Data are reported and presented as median and 95% confidence interval of the median.

We next quantified the relationship between the instantaneous subthreshold DMP magnitude and DAP rate (see Methods) (*58*). The range of subthreshold DMP at which DAP initiated was 4.4-fold larger than the magnitude of the corresponding DAP, and positively correlated with DAP amplitude (Fig. 4e). The large DAP initiation range (Fig. 4e) is similar to that reported for dendrites *in vitro* (*3–8,10*),further supportive of the dendritic origin of our measurements.

Subthreshold DMP strong modulated DAP rates. During SWS, DAP rate had a slowly increasing, sigmoidal dependence on subthreshold DMP (Fig. 4f), with rate increasing over a large dynamic voltage range (see Methods) which often exceeded the magnitude of DAP themselves (Fig. 4f, g, fig. S9c). The firing rate was modulated from nearly 0 Hz at low values of the membrane voltage up to above 100 Hz at the highest membrane voltages (Fig. 4f, g, fig. S9c). Because short-term ion channel dynamics could influence spike initiation properties (*58*), the above calculations were also done for only those DAP separated from all others by at least 50 ms (termed solitary DAP), with only minor differences (fig. S9d-h).

## Dendritic sub- and supra-threshold membrane potential in freely moving rats

The stability of our recordings allowed us to compare dendritic activity and its subthreshold modulation between SWS and during spontaneous, free locomotion (RUN) (Fig. 5a, see Methods). DAP mean rate during RUN (12.8 Hz) was nearly twice as large compared to the mean rate during SWS (6.87 Hz, Fig. 5b), and significantly greater than that of pyramidal neurons (1.99 Hz) or interneurons (4.25 Hz). There was no significant difference in the firing rates between slow and fast runs in DAP. DAP amplitude was slightly smaller during RUN compared to SWS (fig. S6d, e), and rise time and width were slightly longer, though the difference in width was not statistically significant (Fig. 5a, b).

**Fig. 5.**
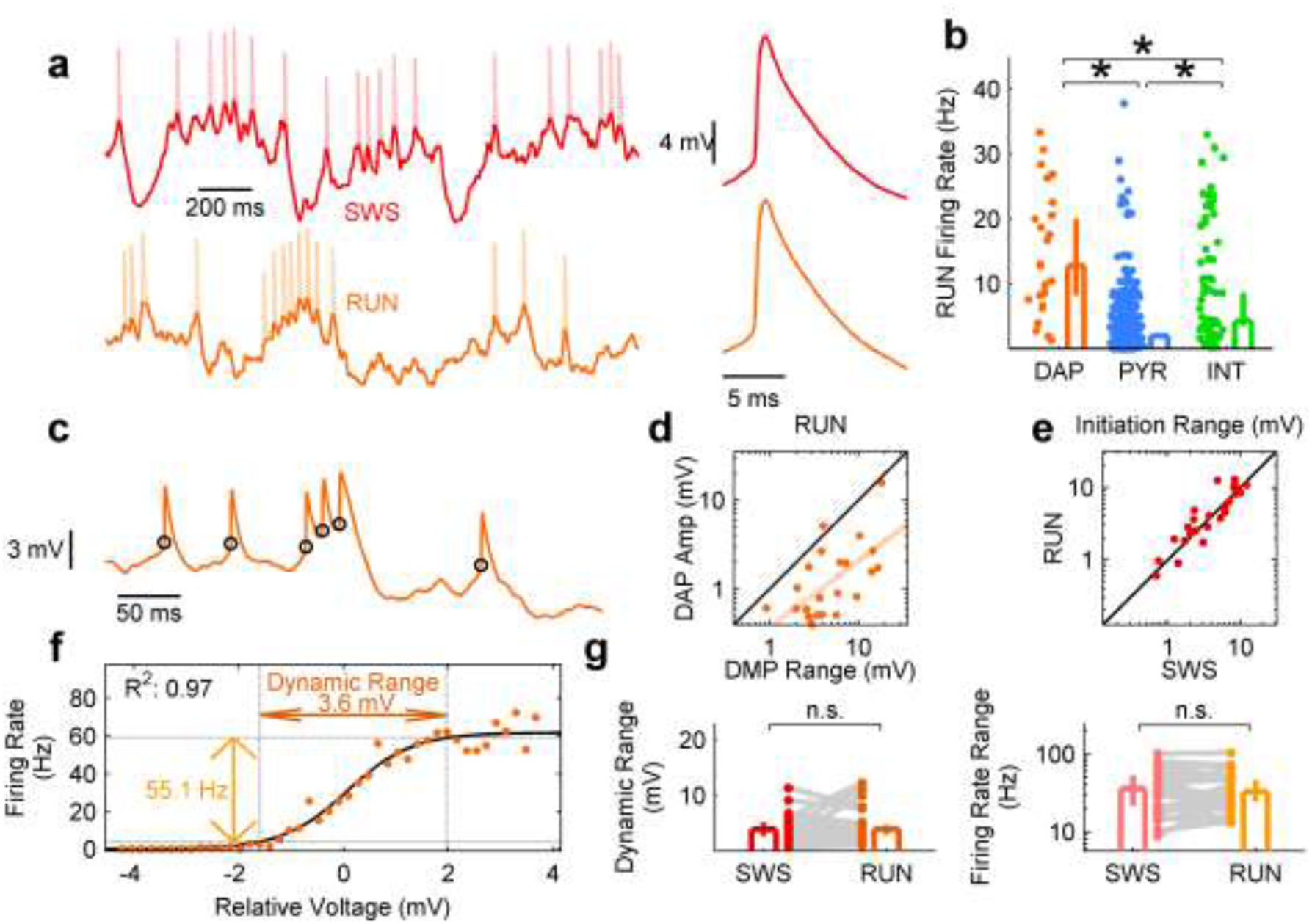
Large subthreshold membrane potential fluctuations modulate DAP rates during RUN. **a**, Sample membrane potentials during SWS (red, left) and locomotion (RUN, orange, right) show similar dynamics and amplitude of both DMP and DAP (middle). **b**, DAP mean firing rate during RUN (12.8, [8.29, 20.0] Hz, n=25 dendrites) was significantly greater than that of pyramidal neurons (1.99, [1.63, 2.37] Hz, n=657 units; p=1.8×10^−5^, Wilcoxon rank-sum test) and interneurons (4.25, [3.57, 8.65] Hz, n=97 units; p=8.3×10^−3^, Wilcoxon rank-sum test) **c**, Sample DMP trace during RUN shows a dynamic initiation range similar to that observed in SWS (Fig. 4e). **d**, DMP range during RUN (3.82, [2.76, 6.17] mV, n=25 dendrites) was significantly larger (p=1.6×10^−5^, Wilcoxon signed-rank test) than the corresponding DAP amplitude (0.82, [0.52, 1.78] mV, n=25 dendrites), and significantly correlated (r=0.68, [0.39, 0.85]; p=2.0x10^−4^, two-sided *t* test). **e**, DAP initiation ranges in SWS (3.67, [2.24, 6.99] mV, n=25 dendrites) and RUN (4.07, [2.51, 8.14] mV, n=25 dendrites) were positively correlated (r=0.91, [0.80, 0.96]; p=3.4×10^−10^, two-sided *t* test) and not significantly different (p=0.46, Wilcoxon signed-rank test). **f**, Sample V-R curve during RUN, which was well-described (Fig. S11) by a sigmoidal logistic function. g, 24 of 25 dendrites had sufficient data to characterize V-R curves in RUN. Left, the dynamic voltage range during RUN (3.90, [2.98, 4.67] mV, n=24 dendrites), was not significantly different (p=0.73, Wilcoxon signed-rank test) from the dynamic range in SWS (3.87, [2.74, 5.03] mV, n=24 dendrites). Right, the firing rate range in RUN (32.5, [24.6, 46.5] Hz, n=24 dendrites) was not significantly different (p=0.86, Wilcoxon signed-rank test) from the range during SWS (35.6, [21.6, 53.1] mV, n=24 dendrites). Data are reported and presented as median and 95% confidence interval of the median. * and n.s. indicates significance or lack of significance, respectively, at the p<0.05 level.

Subthreshold DMP measures were also comparable in SWS and RUN. First, the magnitude of subthreshold DMP was equally large in the two conditions (Fig. 5a, Fig. S10). This is in contrast to the LFP, where fluctuations are severely diminished during locomotion compared to the large fluctuations present in SWS (Fig. S10) (*59*,*60*). The subthreshold DMP magnitude during locomotion was 4.7-fold larger than the corresponding DAP amplitude, and the two were highly correlated (Fig. 5c, d), as in SWS. Second, DAP had a large initiation range in RUN (Fig. 5c, e) that was 5-fold larger than the corresponding DAP amplitude, as in SWS (Fig. S11a). DAP rate during RUN was also modulated by subthreshold DMP in a sigmoidal fashion (Fig. 5f, g, Fig. S11b) over a wide dynamic voltage range, and spanned a large range of firing rates, both of which were as large as in SWS (Fig. 5f, g). These results were similar when performed on solitary DAP during locomotion (fig. S11c-h).

## Modulation of DAP, DMP, and soma by behavior

Do DAP and DMP contain information about instantaneous behavior? Previous studies have shown that somatic spike rates in the posterior parietal cortex (PPC) are modulated by specific types of movements, including forward running, left turns, and right turns (Fig. 6a) (*61, 62*), and have an anticipatory component to their response (*61*). (Fig. 6b, fig. S12a, see Methods). We computed the normalized dispersion, or spread, and depth of modulation of egocentric response maps for parietal DAP rate, subthreshold DMP voltage, and somatic spike rate (Fig. 6b, fig. S12a, see Methods) during free locomotion in a rest box and a random foraging task (fig. S12b, see Methods). The anticipatory component was quantified by finding the time lag corresponding to the best tuning (fig. S12c, see Methods).

**Fig. 6.**
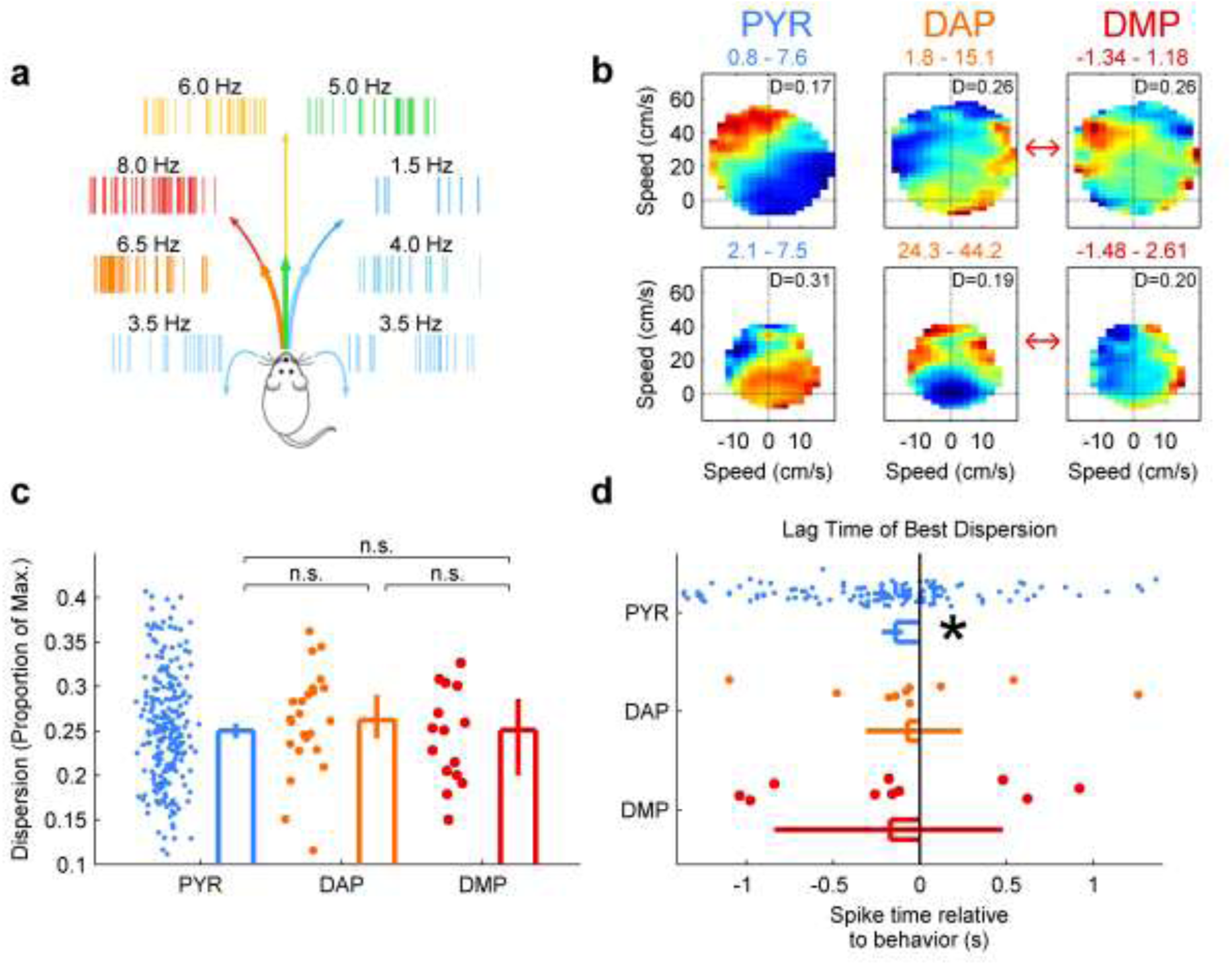
DAP and DMP exhibit egocentric tuning comparable to somatic spikes. **a**, Schematic of egocentric map computation. In this example, the neuron fires maximally (8 Hz, red) during a left hand turn at high velocity, and minimally (1.5 Hz, blue) during a right hand turn at high velocity. **b**, Two sample pyramidal (PYR, left), DAP (middle), and DMP (right) egocentric maps. The minimum and maximum firing rates (mean voltages for DMP) for each map are displayed in the title, and the normalized dispersion D (see Methods, fig. S10a, b) is displayed in the upper-right corner. Red arrows indicate DAP and DMP from the same recording sessions. **c**, The normalized dispersion (see Methods) of pyramidal somata (0.25, [0.24, 0.26], n=245 maps) was not significantly smaller than that of DAP (0.26, [0.24, 0.29], n=24 maps; p=0.4, Wilcoxon rank-sum test) and DMP (0.25, [0.20, 0.30], n=15 maps; p=0.57, Wilcoxon rank-sum test). DAP and DMP dispersions were not significantly different from each other (p=0.33, Wilcoxon rank-sum test). **d**, The time lag corresponding to the optimal tuning for pyramidal somata with significant tuning (−140, [−220, −100] ms, n=146 maps with significant tuning) was significantly different from 0 (p=9.4×10^−7^, Wilcoxon signed-rank test). The same measure for DAP (−70, [−310, 240] ms, n=10 maps with significant tuning) was not different from 0 (p=0.61, Wilcoxon signed-rank test). DMP lag (−170, [−840, 480] ms, n=10 maps with significant tuning) was also not significantly different from 0 (p=0.43; Wilcoxon signed-rank test), and not significantly different from either PYR (p=0.94, Wilcoxon rank-sum test) or DAP (p=0.5, Wilcoxon rank-sum test). Data are reported and presented as median and 95% confidence interval of the median. * indicates significance at the p<0.05 level, and n.s. indicates lack of significance at the p<0.05 level.

There were similar properties of egocentric tuning in pyramidal somata (PYR), DAP, and DMP (Fig. 6c). Normalized dispersion for PYR (0.25) was slightly but not significantly better than that of DAP (0.26, p=0.40, Wilcoxon rank-sum test) and DMP (0.25, p=0.571.4×10^−2^, Wilcoxon rank-sum test), which were also not significantly different from each other (p=0.33, Wilcoxon rank-sum test). The depth of modulation of egocentric maps was comparable between all three groups (fig. S12d), with a similar percentage having a depth of modulation significantly above chance (PYR, 146/245 (60̅, [53, 66]%), DAP, 10/24 (42, [22, 63]%), DMP, 10/15 (67, [38, 88]%)). Coherence showed a different pattern, with DMP having the highest coherence (fig. S12e), but this is likely an artifact because coherence is computed on unsmoothed rate maps, and the continuous DMP signal has more data than spiking point processes. Similar to dispersion, both short-term (fig. S12f) and long-term (fig. S12g) stability for PYR was larger than DAP and DMP, but the differences were not statistically significant.

The time lag corresponding to the optimal depth of modulation for pyramidal neurons with significant tuning was significantly negative(*61*) (−140 ms, p=9.4×10^−7^) (Fig. 6d), indicating a preference to code for future movements. In some instances, somatic spikes were best tuned for behavior several seconds in the future or past (fig. S12h). The optimal lag time for significantly tuned DAP (−70 ms) and DMP (−170 ms) was not significantly different from 0 (DAP, p=0.50; DMP, p=0.43).

Hence, pyramidal neuron somata and dendrites both code for egocentric movement, but with differences that illustrate potential computational principles within a neuron. Pyramidal somatic responses are less dispersed than DAP and DMP responses, even though equivalent percentages of PYR, DAP, and DMP are significantly tuned. The optimal coding occurs at negative time lags, or prospectively, for pyramidal somatic spikes but not for DAP or DMP.

## Conclusions

We directly recorded the membrane potential and sodium spikes of putative neocortical distal dendrites in freely behaving rats without the confounding effects of recent anesthesia or restraint typically used with *in vivo* patch clamp or optical imaging procedures. These measurements were done at sub-millisecond temporal resolution and were stable for at least one hour and up to four days. DAP kinetics were similar to those seen *in vitro* and in anesthetized animals (*3–8,10*). These recordings revealed a number of interesting properties of dendritic activity. First, DAP fired at very high rates *in vivo*, far greater than somatic rates. Second, DAP were accompanied by large subthreshold DMP fluctuations, the magnitudes of which were always larger than the DAP amplitude. Third, this was present not only during SWS but also during free locomotion. Fourth, DAP rates varied by an order of magnitude as a function of the subthreshold DMP in a graded fashion during both SWS and movement, suggesting a large dynamic range. Fifth, unlike previous reports (*23,24*), our measurements from PPC showed substantial differences between DAP, DMP and somatic spikes in both quality and temporal dynamics.

## Discussion

DAP have long been hypothesized to endow neurons with greater computational power by turning dendritic branches into computational subunits with branch-specific plasticity (*9,18–21*). Dendrites support initiation and propagation of action potentials *in vitro* via voltage-gated sodium, potassium, and calcium channels (*3–8,10*). It has been unclear if the conditions for generating DAP exist *in vivo* during natural behavior. Patch-clamp recordings in head fixed mice suggest that proximal dendrites in visual cortex support spikes, though a majority of them were bAP while some were DAP (*24*). In contrast, our recordings are putatively from electrotonically distal-most dendrites which generate dendritic spikes locally, and where somatically generated bAP are mostly absent (*2,6,7,10,16*). These recordings were stable for hours or days at a time during free locomotion with no evident damage to the dendrite.

We hypothesize that our tetrodes are in a “glial-assisted” configuration similar to in-cell or quasi-intracellular recordings (*29,30,33,34*). In this model the requisite high seal resistance comes from glial encapsulation (*37,38*) trapping a segment of a thin dendrite between the four electrode tips and the glial cells, forming a stable configuration providing high quality, positive polarity sub- and supra-threshold signals. Our immunohistochemical labeling and *in-vivo* impedance spectroscopy measurements support this model, and the negative correlation between the subthreshold DMP and LFP is consistent with the whole cell results from the soma (*57*). The encapsulation process might be facilitated by the fine-scale geometry of our gold-plated tetrode tips. However, the tetrodes have to be positioned close enough to a neuron to trap the dendrite without puncturing it. Indeed, our current success rate of dendritic recordings is quite low but comparable to the success rate of somatic whole cell recordings during natural behavior (*44,63,64*).

Several signal properties, including dendritic spike rise time and width, firing rate, and short-term plasticity, are consistent with the hypothesis that our measurements are from the electrotonically distal-most dendrites of pyramidal neurons and the dendritic spikes are generated locally in the dendrites, not back-propagated from the soma. The high DAP rates are unlikely to come from damage to the neuron due to stable DAP properties over hours. Spike properties such as width and complex spiking indicate that our data are from the pyramidal neurons but not inhibitory interneurons. This may be because less than 20% of neurons are interneurons and their dendritic arbor is far smaller than the pyramidal neurons', thus reducing the chances of encountering interneurons. The dendritic spike properties were unimodal, with low variability and no refractory violations, indicating a single dendritic source. While many dendrites could be potentially trapped between the tetrode tips, very few would remain intact, reducing the success rate and chances of multiple source. This is consistent with recording two order of magnitude less (than expected) extracellular cells using tetrodes (*40*).

Because these rates greatly exceed those of pyramidal units in the same brain region, a large amount of activity and information processing in a neuron may be occurring in the dendrites without being read out in the postsynaptic soma, consistent with *in vitro* studies showing decoupling of somatic and dendritic compartments (*2,6,7,10,13,16*). The results of DAP computations could travel to the rest of the network via second messengers and presynaptic mechanisms. The semi-independence of distal dendrites suggests that NMDA-dependent plasticity, that crucially depends on depolarization, could be induced in only a specific dendritic branch, allowing for more input-specific plasticity and clustering of synapses with similar information (*9,13,18–21*) and greatly expanding the computational capacity of a single neuron.

DAP firing rates are modulated in a graded fashion by the subthreshold DMP. This endows the dendrites with an analog code, defined by the depolarization level of the dendrite, reminiscent of the sigmoidal response profile of the hidden layer of artificial neural networks (*18,19*). The digital coding, carried out by DAP, coexists with the subthreshold modulation, indicating a hybrid analog-digital code in the dendrites. This large subthreshold modulation range (~5 mV average, ~20 mV max) is consistent with but greater than the recent results from the somata of hippocampal CA1 neurons (*43, 44, 64*), entorhinal stellate cells (*65*), and barrel cortex neurons (*66*). The true subthreshold voltage range in our case is likely even greater, since the magnitude of signals obtained through our measurement technique would be smaller than the true range, and influenced by the quality of the glial seal.

Somatic spikes in PPC are modulated by movement in an egocentric reference frame (*62*) in an anticipatory fashion during free behavior (*61*). DAP and DMP also exhibit egocentric modulation, but are more disperse and do not show significant anticipatory behavior. Our findings suggest a computational framework in which individual cortical neurons take information about the current state of the world, present in the dendrites, and form an anticipatory, predictive response at the soma, a computation similar to that performed by a Kalman Filter (*67*) or Hidden Markov Model (*68*). Unlike network models of sequence learning, each individual neuron may behave like a feed-forward circuit that performs predictive coding based on non-predictive inputs arriving from many dendritic branches (*20,69–72*). The intermediate integration step performed by the dendrites is likely a crucial one in neurons with extensive dendritic trees, to allow inputs at distal tufts to be integrated in the somatic response (*6*).

These results demonstrate that dendrites generate far more spikes than the soma. This has important implications in many fields. The total energy consumption in neural tissue, measured in fMRI studies, could be dominated by the dendritic spikes(*73*). Correlations between neuronal somatic spikes may be weak (*74*) due to the larger number of dendritic spikes with only a fraction generating somatic spikes. Thus, the abundance of dendritic spikes *in vivo* could profoundly alter synaptic plasticity and facilitate clustering of synapses on dendritic branches, thus altering the nature of neural code and memory.

## Materials and Methods

### Recording Procedure

#### Subjects

Data were obtained from 9 singly housed adult male Long-Evans rats (350-425g at the time of surgery). Animals were trained to perform spatial exploration tasks non-central to the present study and described previously (*35,36*). The animals were water restricted (minimum of 30mL/day) in order to increase motivation to perform tasks, and received sugar water or solid cereal rewards during spatial exploration tasks. Further, they were food restricted (minimum of 15g/day) to maintain a stable body-weight and increase motivation to perform in the random foraging task (see below). All experimental procedures were approved by the UCLA Chancellor's Animal Research Committee and in accordance with NIH-approved protocols.

#### Surgery and *in vivo* electrophysiology

All methods were analogous to procedures described previously (*35, 36*). Rats with satisfactory behavioral performance on spatial exploration tasks were anesthetized using isoflurane and implanted with custom-made hyperdrives with 22 independently adjustable tetrodes. All implanted tetrodes were constructed in-house from ~13 μm-diameter NiChrome wire coated with ~2 μm-thick polyimide insulation, according to previously-reported techniques(*35, 36*). After the tetrodes were cut, the tips were electroplated with a gold particle solution containing 75% gold plating solution and 25% multiwalled carbon nanotube solution. A current of 1 μA was passed for 2 seconds with the electrode tip as the anode. This was repeated until the impedance at 1 kHz decreased below 250 kΩ for each electrode of every tetrode. Additional checks ensured that the four channels of a tetrode were not shorted.

4 rats were implanted with drives targeting right prefrontal cortex (1.2–2.5 mm lateral, 2.0-3.9 mm anterior to bregma) and right posterior parietal cortex (1.8-3.5 mm lateral, 3.5–4.1 mm posterior to bregma). 5 additional rats were implanted with drives targeting both left and right parietal cortex. After recovery from surgery, all tetrodes were slowly lowered through cortex over a span of several days to months. Subsets of tetrodes were advanced daily, typically ~70 μm and rarely more than 140 μm in one day. Before tetrodes were adjusted each day, all continuous signals were visually and aurally screened for intracellular-like signatures, consisting of large-amplitude fluctuations and broad, large amplitude spikes of reverse polarity compared to normal extracellular units. While a tetrode recorded a dendritic signal, no other tetrodes in the same hemisphere were adjusted. DAP signals always manifested overnight while all tetrodes were stationary. 25 independent dendritic sources were recorded across these 9 rats, while no intracellular-like signals manifested in 14 other rats implanted using similar procedures.

Signals were recorded using a Neuralynx data acquisition system at a sampling rate of 32 kHz from 7 rats and 40 kHz from 2 rats; all signals were initially digitally filtered through a 32 tap low-pass FIR filter with a cutoff frequency of 9000 Hz and a high-pass DC offset filter with a cutoff frequency of 0.1 Hz. Data from both brain regions and both hemispheres were pooled together for Figures 1–5 and figures S1–2 and S5–12, as no systematic differences existed between any of these regions. Data from both parietal hemispheres from 5 rats recorded in the rest box and while performing the random foraging task (see below) were pooled together for Figure 6 and figure S12.

### Behavior

#### Position tracking

For 5 rats, the animal’s position was measured using an overhead camera that detected the position of colored LEDs (0.5 cm x 0.5 cm each) placed on top of the electrode assembly on the head, and sampled at an average of 55 Hz at a resolution of 640x480 pixels. This data was used to compute instantaneous position, running speed and movement direction.

#### Rest box

The majority of data were recorded when rats were left to freely behave in a “rest box” made of plastic with an open top (60×40 cm, 103 cm high). A small cloth was placed in the box for rats to rest on, but no other salient cues were present on any surface of the box, and the box was cleaned between recording sessions to eliminate odors. During recording, rats were totally unrestrained and free to groom, sleep, and move around; hence we refer to behavior in this condition and random foraging (see below) as “free locomotion”. This configuration allowed us to characterize the properties of the same DMP during SWS and RUN. Recording sessions lasted approximately one hour, and between one and three recording sessions were performed per day, interspersed with task sessions referenced above. Position data was recorded for 5 rats in this condition, yielding a total of 8 DAP across a total of 16 recording sessions.

#### Random foraging

Data from one rat was also obtained during a task where the rat foraged for randomly dispersed food rewards in an open field 100×100 cm with 50 cm high walls, each with a distinct visual cue. Each session lasted approximately 20 minutes. 4 DAP were recorded across a total of 8 recording sessions in this condition. There were no notable differences between data from the Rest Box and Random Foraging conditions, and so were combined for analysis for Figures 5, 6, and figures S10, S11,and S12.

#### Behavioral state identification

Spectral features of the LFP were used to estimate behavioral state. The 32 (40) kHz continuous traces were downsampled by a factor of 8 to 4 (5) kHz and digitally filtered with a zero-phase second order Butterworth bandpass filter with cutoff frequencies of 0.5 and 500 Hz. Using the built-in MATLAB function spectrogram(), the power spectral density of the LFP was estimated using five-second periods shifted by one second, yielding a spectrogram with a frequency resolution of 0.4 Hz and a temporal resolution of 1 second. This spectrogram was smoothed by 2-dimensional convolution with a Gaussian kernel of standard deviation 0.4 Hz and 1 second.

Using the resulting spectrogram, rat behavior was manually classified into two categories: slow wave sleep (SWS) and running (RUN). Time periods corresponding to whisker twitching, sleep spindles, handling by experimenters, or other movement artifacts were discarded. SWS was defined as time periods with elevated power in the 1.5–5 Hz (delta) band, reflecting large amplitude cortical up and down states (UDS). Remaining time periods were defined as RUN, long segments at least 20 s long, typically lasting several minutes. These segments included fast movements, slow movements and brief periods of relative quiescence. When available, these classifications were confirmed by tetrodes located in the CA1 region of the hippocampus showing elevated power in the 6–12 Hz theta band, a typical signature of movement, or directly with video recording.

### Extracellular electrophysiology and spike sorting

#### Extracellular unit classification

Spike extraction, spike sorting and single unit classification were done offline using custom software and according to methods described previously (*35,36*). Extracellular waveforms were extracted from the LFP filtered between 300 and 9000 Hz with a zero-phase seventh order Butterworth bandpass filter. Peaks with values above an adaptive threshold based on the magnitude of the noise in the signal (typically >40 μV) were identified as putative somatic action potentials.

Because we compare firing rate properties between extracellular units and DAP, we used the spike waveform to classify the putative neuron type of extracellular units. After manual spike sorting to assign spikes to isolated units, the average spike waveform was estimated as the mean across all spikes within a cluster. The width at half-maximum and time from spike peak to trough were then computed for each cluster. Only units from neocortex with a mean firing rate above 0.05 Hz in either SWS or RUN, as well as a half-width below 0.45 ms were included in all analyses. Units were classified as either pyramidal neurons or interneurons based on a combination of these measures. Units satisfying 1.6*W + P > 0.95 were identified as putative pyramidal neurons, and units satisfying 1.6*W + P <= 0.95 were identified as putative interneurons, where W is the width at half-maximum and P is the time from spike peak to the trough immediately after the peak. The parameters defining this equation were chosen to achieve maximal separation between two clusters in the width vs peak-to-trough space, and are consistent with previous studies (*45*).

#### Complex spike index (CSI)

All CSI data presented were computed only using spikes in SWS to eliminate potential behavioral bias. For all pairs of adjacent spikes with spike time T_n_ and amplitude A_n_ and belonging to a single unit, the inter-spike-interval (ISI) was defined as

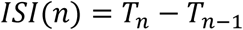

and the adaptation index (ADI) was defined as the ratio

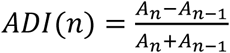

The CSI was then computed as

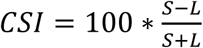

where S is the number of spike pairs with ISI < 20 ms and ADI < 0, and L is the number of spike pairs with ISI < 20 ms and ADI >=0. Any other spike property can be substituted for amplitude to quantify the degree of change with repeated activation. Since the rising phase of the derivative of intracellular somatic spikes corresponds to the rising phase of extracellular spikes (*40*), the DAP CSI reported in Figure 3e and 3f is computed for the peak value of the first temporal derivative. CSI computed for DAP rise time and width are reported as the negative of the above formula, as rise time and width were observed to increase with repeated activation, rather than decrease.

#### MUA estimation

All MUA data presented were computed only using spikes in SWS. For a given tetrode, only detected spikes with a half-width between 0.05 and 0.45 ms and a peak-to-trough time between 0.3 and 1.2 ms were included in MUA calculations. This eliminates contamination by spurious events. The MUA amplitude for a given tetrode was computed as the mean of all spike amplitudes across all four tetrode channels, and the rate was computed as the number of spikes divided by the amount of time spent in SWS. MUA rate estimated thusly is not independent of MUA amplitude, as spikes with an amplitude below the adaptive detection threshold will not contribute to the MUA rate.

#### Amplitude Variation

Given the amplitude of a spike (Extracellular or DAP) on all four channels of a tetrode *A*_1_, *A*_2_, *A*_3_, *A*_4_ and the mean amplitude

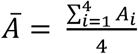

the amplitude variation is defined as

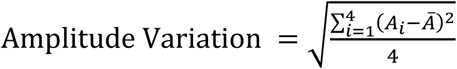

For recording sessions prior to (PRE) and following (POST) dendritic recording, amplitude variation was computed over the amplitude of all spikes from a single extracellularly-measured unit. During dendritic recording, amplitude RMS was computed over the amplitude of all spikes from a single DAP.

### Dendritic detection and quantification

#### DAP Detection

Detection of putative DAP was performed on the individual tetrode channel with the largest amplitude-fluctuations. The continuous trace was filtered below 1500 Hz with a zero-phase second order Butterworth lowpass filter. Positive peaks in the first temporal derivative were identified as putative DAP, and the peak value of the second derivative immediately before the first derivative was recorded for each. Using these times, analogous values were obtained on each of the other tetrode channels. DAP were manually separated from noise offline using custom MATLAB scripts based on the first and second derivative peaks on all four channels (Fig. S1).

#### DAP Quantification

The initiation time of a DAP was defined as the time the first derivative crosses above 25% of its peak value immediately before the peak (Figure 1d). The end of the rising phase of the DAP was defined as the time the first derivative falls below 25% its peak value. DAP rise time and amplitude were defined as the time difference and voltage difference, respectively, between these two points. Due to the large range of signal amplitudes across recordings, we did not establish a static threshold of *dV/dt* to detect DAP in order to eliminate biases. The DAP width was defined as the time difference between, half-amplitude crossing points identified up to 3 ms before DAP peak and 50 ms after DAP peak (Figure 1d). If no such points were identified for a particular DAP (<5% of all spikes), the width was excluded from further analysis. The estimated width of the extracellular DAP reported in Figure 3d was computed in a similar fashion, using the first temporal derivative, filtered between 300 and 9000 Hz, with a zero-phase seventh order Butterworth bandpass filter, as done for all extracellular units. For each DAP, the mean values reported for amplitude, rise time, width, and extracellular width were computed from the mean of all spikes separated from all other spikes by at least 50 ms.

#### Duration of Recording

The duration of each DMP recording was defined as the time difference between the end of the last session and the beginning of the first session containing intracellular signatures on a given tetrode. Hence, the reported hold times are likely underestimates of the true DMP hold time. A signal recorded on the same tetrode on consecutive days was designated to be the same DMP based on the similarities of DAP waveform and firing rate, and unchanged tetrode depth.

#### Peak Rate and Inter-spike Interval Calculations

The shortest ISI reported in figure S5 was calculated as the lowest 5% of the ISI histogram. The peak rate reported in figure S6 was defined as the reciprocal of the lowest 5% of the ISI histogram.

#### Immunohistochemistry

After terminal anesthesia by IP injection of 40 mg/kg pentobarbital, the brain was immediately removed and immersion-fixed in 4% formalin for 72 hours, washed and cryoprotected in a 30% sucrose in 0.1M phosphate buffer (PBS). The brain was embedded in OCT compound (Sakura) and frozen on dry ice. 30-μm thick coronal sections were cut on a cryostat and the sections stored at 4°C in PBS with 0.06% NaN_3_ until processing.

Specific sections with evidence of tetrode tracks were selected for processing. The sections were washed (3x10 minutes) in 0.1M phosphate buffer, 0.9% NaCl pH 7.3 (PBS) and incubated at room temperature (RT) in a blocking buffer of 5% normal donkey serum (NDS)/0.3% Triton-X for 1 hour. Primary antibodies, applied with 2% NDS/0.3% Triton-X overnight at RT, were: rabbit anti-ionized calcium-binding adapter molecule 1 (Iba1) (1:500, Wako); mouse anti-glial fibrillary acidic protein (GFAP) (1:50, Thermo Fisher Scientific); chicken anti-microtubule-associated protein 2 (MAP2) (1:500, Phosphosolutions). Sections were then rinsed (3x10 minutes) in PBS and incubated in a mixture of secondary antibodies for 2 hours at RT. Secondary antibodies used were donkey anti-rabbit rhodamine-red X (1:250, Jackson Immuno) to visualize Iba1, goat anti-mouse Alexa 488 (1:250, Invitrogen) to visualize GFAP and donkey anti-chicken biotin (1:500, Jackson) to visualize MAP2. The sections were rinsed (4x15 minutes) in buffer and then stained with streptavidin-Alexa 647 (4g/ml) for 1 hour. After a final rinse in buffer (4x15 minutes), the sections were mounted on Superfrost plus slides and allowed to dry briefly.

The sections were coverslipped with ProLong Gold anti-fade reagent, and imaged using scanning confocal laser microscopy. Maximum projection and semi-transparent 3D image stacks were prepared using ImageJ software, using green, cyan, and red look-up tables for GFAP, Iba1, and MAP2, respectively. The dendrite indicated in figure S3a, bottom and Movie S2 was highlighted in magenta using a customized script in MATLAB.

#### Impedance Spectroscopy

As a confirmation of the glial sheath hypothesis described in figure S3a and S3b, we measured the impedance of electrodes *in vivo* during dendritic recording and compared that to electrodes that were recording local field potential (figure S3c, d).While the rat rested in the rest box, the impedance of all channels on 7 tetrodes was measured at 20, 50, 100, 200, 500, 1000 and 2000 Hz, using the same Neuralynx hardware used to monitor impedance during electroplating (see above).

The impedance thus measured is typically modeled as a combination of several components. First, the electrode-electrolyte interface is modeled as a constant phase element (CPE) *Z_el_*. The impedance of a CPE is expressed in the following form:

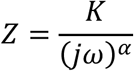

Here *K* represents the overall magnitude of the impedance, *j* is the imaginary number (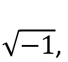) is the frequency, and *α* is constrained between 0 and 1. As *α* approaches 0, the behavior of the CPE approaches that of a pure resistor. As *α* approaches 1, the behavior approaches that of a pure capacitor.

The second component of the modeled impedance is the impedance of the surrounding glial tissue between the electrode and ground. There are two such pathways, one directly through the surrounding glial cells, which is modeled as a second CPE *Z_g_*, and one through the space between glial cells, modeled as a frequency-independent resistor *R_g_*.

For each electrode, the 5 parameters (*K_el_*, *α_el_*, *K_g_*, *α_g_*, *R_g_*) were estimated by minimizing the following function:

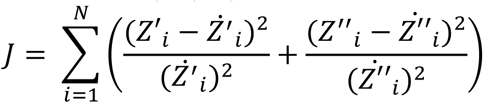

where *i* indicates the different frequencies, N represents the total number of frequencies, *Z*′_*i*_ represents the real part of the measured impedance at the specific frequency, *Ż′_i_* represents the corresponding model estimate, and *Z″_i_*and *Ż″_i_* represent similar values for the imaginary part of the measured impedance. Parameters were estimated using the built-in patternsearch() function in MATLAB. Different sets of initial values were tested and the results producing the minimum error were selected, thus improving the chances of obtaining the global minimum.

All 5 parameters were unconstrained during fitting, but a systematic relationship emerged between the pairs of parameters specifying the CPE elements (*K_el_* related to *α_el_*, *K_g_* related to *α_g_*). A thorough investigation of this relationship is beyond the scope of the current study, so only the *K* values for each CPE are reported.

Electrode wire axial resistance and leak capacitance (not depicted in figure S3b) are typically orders of magnitude smaller than the impedances shown, as is *R_ax_*, the axial resistance within the dendrite. *R_amp_*, the resistance across the amplifier used to record signals, is typically several orders of magnitude larger than the impedances shown. For these reasons, these parameters were ignored in all analyses.

#### Compartmental Model

We simulated a layer ν pyramidal neuron by modifying a previously-published model (*53*) implemented in NEURON version 7.2 (*54*). All membrane properties and channel densities were identical to those previously used with exceptions described below. The simulation was run with a time step of 0.025 ms, and 1000 ms were simulated. Each segment was simulated with at least 6 compartments.

Stimulation to the neuron was given using in-vivo like conductances (*55*) governed by an Ornstein-Uhlenbeck process (*56*). Briefly, an excitatory and inhibitory current are modulated by conductances g_e_ and g_i_ which have mean values g_e0_ and g_i0_ and are perturbed by Gaussian white noise scaled by coefficients D_e_ and D_i_. To generate DAP, we selected a single compartment in a distal dendrite to inject this input. bAP were simulated by injecting current into the soma.

Voltage traces from each segment of the model were written to file and then analyzed in MATLAB after being passed through a 32 tap low-pass FIR filter with a cutoff frequency of 9000 Hz, as is done with the experimental data. Rise time and width reported in Fig. S7 were computed from the mean waveforms triggered on the time of spikes in the compartment in which stimulation was injected.

Parameters differing from those previously published (*53*) are as follows: E_passive_ (passive reversal potential) = −50 mV; P_HVA,dend_ (high voltage-activated Ca^2^+ channel permeability) = 0.00016 μm/s; G_Na_bar_ (fast sodium channel peak conductance) = min(1000, B*400 pS/μm^2^, where B = 1/diameter^2^ for dendrites farther than 650 μm from the soma.

Further, the conductance parameters for dendritic stimulation are as follows: g_e0_ = 0.028 μS; g_i0_ = 0.0573 μS; D_e_ = 0.02 μS; D_i_ = 0.0066 μS; For somatic stimulation: g_e0_ = 0.007 μS; g_i0_ = 0.0573 μS; D_e_ = 0.007 μS; D_i_ = 0.0066 μS.

### Subthreshold Responses

#### Subthreshold Magnitude Estimation

To eliminate the possible influence of long timescale fluctuations in signal properties, all subthreshold analyses were performed on segments of data approximately 5 minutes in duration. To eliminate possible influences of refractoriness or biasing of the voltage due to the large DAP amplitude, the subthreshold trace was constructed by eliminating all data in the original voltage trace 4 ms after each DAP initiation point. The membrane potential range reported in Figures 4b-c, 5d, and figure S10d was defined as the difference between the 5^th^ and 95^th^ percentile of the distribution of voltages in the subthreshold trace.

#### Initiation Range

The DAP initiation voltage reported in Figures 4d, 5e, and figures S9d and S11a, c, e was defined as the subthreshold voltage at the time of DAP initiation. The initiation range for a given DAP was computed as the difference between the 5^th^ and 95^th^ percentile of the distribution of DAP initiation voltages.

#### Voltage-Rate (V-R) Curve

For each trace, the subthreshold voltage was split into 100 equally-spaced bins. The firing rate in each voltage bin was computed by dividing the number of DAP initiated in each voltage bin by the amount of time the DMP sampled the corresponding voltage bin. Voltage bins with less than 300 ms of data were excluded from analysis to eliminate artifacts. Using the built-in MATLAB function nlinfit(), a logistic function of the following form:

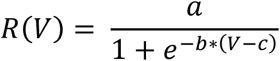

was then fit to the resulting data points, where a, b, and c are estimated constants.

The “dynamic voltage range” reported in Figures 4f, 5g, and figures S9e and S11f was defined as the difference between the voltages at which the V-R curve crossed 5% and 95% of the maximum firing rate of the V-R curve. The “firing rate range” reported in Figures 4f, 5g, and figures S9f and S11g was defined as the difference in firing rate between these two points. Two DAP sources did not have enough spikes in RUN to generate a V-R curve, and these were excluded from analysis in Figure 5g and figure S11f-h.

#### Solitary DAP Analysis

To isolate the influence of a DAP on subsequent DAP and DMP via adaptation or short-term plasticity, a second subthreshold trace was constructed as above but with 50 ms of data excluded after each DAP initiation point. All subthreshold analyses above were repeated on these DAP isolated data for figures S9 and S11.

### Egocentric Responses

#### Position Tracking

Position originally sampled at an average of 55 Hz was interpolated to a uniform 50 Hz. To eliminate abrupt position changes due to transient LED occlusion, position was first smoothed using a median filter of 250 ms width. Position was then further smoothed using a 15-point moving mean filter, and then up-sampled by linear interpolation at all data points including the occluded data to 100 Hz. Instantaneous heading direction H(t) was calculated as done previously (*61*):

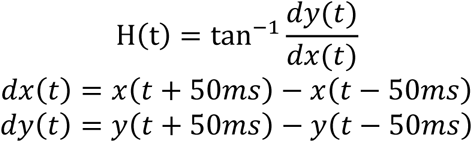

#### Construction of Egocentric Rate Maps

Movement in an egocentric reference frame was computed frame-by-frame by calculating the difference in position and heading direction between the start and end of a moving 100 ms time window.

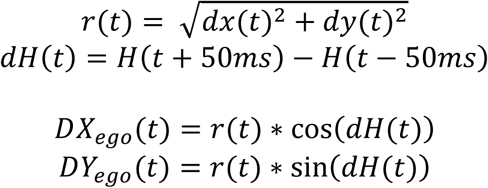

Two-dimensional egocentric movement (*DX_ego_*(*t*) and *DY_ego_*(*t*)) was then down-sampled to 50 Hz, binned using 0.25 cm bins in egocentric space, and smoothed by 2-dimensional convolution with a Gaussian kernel of standard deviation 0.25 cm in each dimension. Egocentric firing rate maps for pyramidal somata and DAP were generated by dividing the number of spikes occurring in each egocentric bin by the total time spent in the corresponding bin. Egocentric voltage maps for DMP were generated by calculating the mean value of the voltage at all times a bin was occupied. Bins occupied less than 250 ms for every 20 minutes of recording were discarded. Maps constructed with 0.25 cm bins were used for all figures, and all statistics (below) were calculated from maps constructed with 0.15 cm bins as done previously (*61*). For figures, displacement (cm) was converted to speed (cm/s) by dividing by 100 ms.

For evaluation of anticipatory responses and to obtain control data, maps were constructed as above for different time lags of spikes (or subthreshold voltage) with respect to behavior. For a given unit, the time of all spikes (or voltage measurements) were circularly shifted by an amount between ±20 seconds, with a resolution of 20 ms. Maps were constructed as above from the resulting data, and measures of selectivity (defined below) were computed for all time lags, smoothed with a Gaussian smoothing kernel of standard deviation 60 ms. The time lag with the highest depth of modulation (ΔS/S, see below) was identified as the peak lag time. For all figures and analyses except figure S12h, the peak lag time was restricted to be between ±1.4 seconds.

#### Depth of Modulation

Traditional measures such as information content or sparsity assume that the data takes strictly positive values (which is not valid for the change in membrane potential) and are not shift-invariant. Hence, to compare selectivity of DMP, DAP and somatic spikes, we first computed the standard deviation of response maps. This measure allowed the quantification of subthreshold DMP maps, which take negative values. The standard deviation S was expressed as a percentage change from baseline, resulting in a ΔS/S signal which is both shift- and scale-invariant, akin to the ΔF/F signal used in calcium imaging. Measured thus, maps with poor tuning will be uniform and have a low ΔS/S, while maps with high values concentrated in a few pixels will have a high ΔS/S. This allows comparison of units with disparate firing rates as well as allow comparison of maps representing firing rates (PYR, DAP) with those representing voltage (DMP). ΔS/S was calculated from smoothed rate maps with 0.15 cm bins.

For each unit, the baseline was calculated using a resampling procedure. As above, the time of all spikes (or voltage measurements) were circularly shifted by an amount between ±20 seconds in either the negative or forward direction, with a resolution of 20 ms. The standard deviations of the resulting maps with lags between ±1.4–20 seconds formed a null distribution, and the median of this distribution was taken as the baseline standard deviation S_0_. The quantity ΔS/S was thus quantified as:

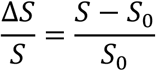

Units were identified as having significant tuning if the peak ΔS/S value within ±1.4 seconds was greater than the ΔS/S at any other time lag. All other measures described below were evaluated only at the time lag of maximal ΔS/S.

#### Dispersion

Depth of modulation quantifies the variation in pixels in a response map, but contains no information about how concentrated, or clustered, pixels with similar values are. We thus computed the “smoothness” of response maps by calculating normalized dispersion. Dispersion was computed on smoothed response maps with 0.15 cm bins (see above). Dispersion was calculated as the mean distance (in centimeters) between the 10% of pixels with the highest values. The distance between such highly active pixels is influenced by the range of running speeds attained by the rat, which can vary across sessions. Hence we normalized the dispersion by the range of displacements in the forward and backward directions, i.e. the distance between the most negative and positive y values in the rate map (Figure 6b).

#### Coherence

Egocentric map coherence was calculated by evaluating the correlation coefficient between each pixel of a map and the mean of all surrounding pixels^45^. Pixels with undefined values were excluded from this analysis. Coherence was computed on unsmoothed response maps with 0.15 cm bins and Fisher z-transformed.

#### Short-term Stability

Short-term stability was evaluated as the Fisher z-transformed correlation of response maps constructed with data from every alternate minute (minutes 0–1, 2–3, etc. compared with minutes 1–2, 3–4, etc.).

#### Long-term Stability

Long-term stability was evaluated as the Fisher z-transformed correlation coefficient of maps constructed with data from the first and second halves of a given recording session (For a 20 minute session, minutes 0–10 compared with minutes 10–20).

#### Number of Dendrites

Multiple recording sessions for a dendrite were kept separate for egocentric analyses. Thus the dendritic sample size for egocentric comparisons is the number of sessions across all dendrites recorded with position data. In total, DMP were recorded with position data in 15 recording sessions across 5 different tetrodes. In several of these sessions, multiple independent DAP sources were present with a single DMP source, for a total of 24 DAP maps.

### Statistics

#### Significance Tests

All analyses were done offline using custom-written codes in MATLAB. Due to the relatively small dendritic sample size and potential non-Gaussian distribution of measures, we employed non-parametric tests and resampling statistics to assess statistical significance. Significance between unpaired data was assessed using the Wilcoxon rank-sum test. Significance between paired data was assessed using the Wilcoxon signed-rank test. These tests make few assumptions about the distributions of data being tested and are robust to non-equal sample sizes or non-Gaussian nature of data. Correlation coefficients and their related significance were calculated using the built-in corrcoef() function in MATLAB, which calculates a two-sided *t* statistic to assess significance.

#### Confidence Intervals

Unless otherwise stated, all values are reported as median [95% confidence interval], in the form M [L, U], with M representing the median and L and U representing the lower and upper bounds, respectively, of the 95% confidence interval. Confidence intervals were estimated using resampling statistics to allow analysis of non-Gaussian distributions. Briefly, a surrogate population was constructed by drawing with replacement from the original distribution and the median of the resulting distribution recorded. This was repeated 100,000 times to form a distribution of the estimated median. The cutoff values of the 2.5^th^ and 97.5^th^ percentile of the estimated distribution were designated as the 95% confidence interval of the original population.

#### Validation of Detection Algorithm and Generation of Surrogate High Rate Data

The fidelity of the detection algorithm was validated by constructing surrogate data sets with known spike times. A typical DAP waveform normalized to an amplitude of 1 mV was convolved with impulses spaced at varying distances, and Gaussian noise with standard deviation of 0.1 mV was added. The detection algorithm was run on the resulting trace. The minimal interval between spikes the algorithm could detect was 0.4 ms. The noise level plotted in figure S5e was computed as the 99.99^th^ percentile of the distribution of peaks in the derivative of the surrogate signal, and serves as a visual guide.

To verify that the high DAP firing rates were not a product of multiple, independent sources being pooled together, surrogate data was simulated and quantified in figure S5e-g. Inter-spike intervals (ISI) were generated from a gamma distribution with shape parameter 1.75 and an appropriate scale parameter to generate a mean ISI of *1/f*, where *f* is the desired firing rate. The shape parameter of 1.75 was chosen to eliminate the majority of ISI less than 2 ms and to fit the observed ISI distributions of DAP. Few remaining ISI <3 ms (<1% of all data) were discarded. Spike times were then generated by taking the cumulative sum of the intervals. Impulses were scaled by a random amplitude drawn from the distribution of amplitudes from a sample DAP (mean 1.5 mV), and then convolved with a canonical DAP waveform with peak value 1. Finally, Gaussian noise with standard deviation of 0.05 mV was added. The detection method described above was then applied to the resulting traces to compute the histograms shown in figure S5f and g.

**Fig. S1.**
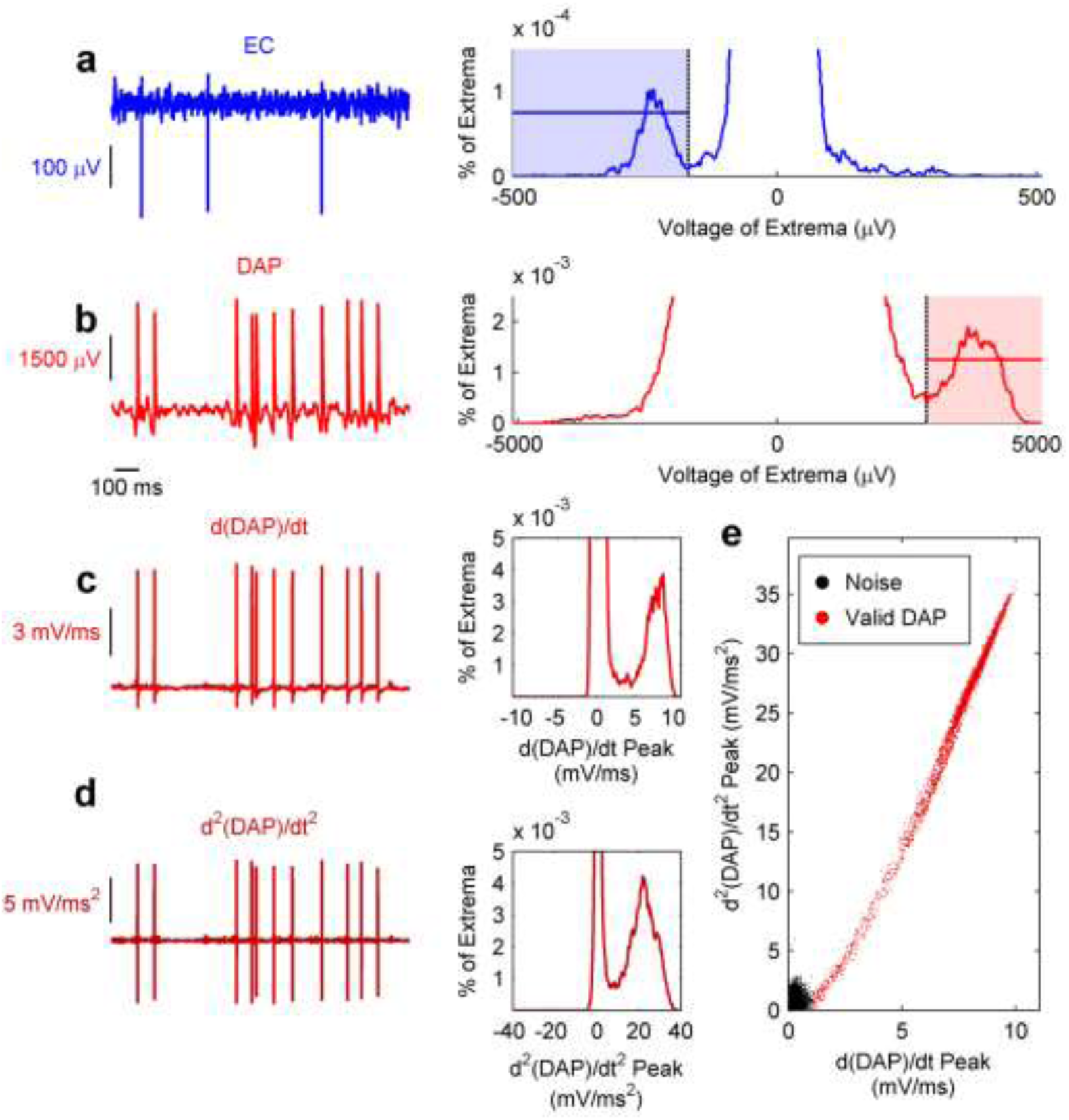
Distributions of somatic spikes, DAP and its derivatives. **a**, Left, example segment of an LFP trace filtered between 20 and 9000 Hz showing negative-polarity spikes. Right, histogram of all extrema (maxima and minima) detected in the entire LFP trace, showing a prominent cluster of peaks around −240 μV (shaded) and no prominent cluster of positive extrema. The y-axis is limited to cut off the large number of peaks in the noise cloud. **b**, Left, example segment of an MP trace similarly filtered showing positive-polarity spikes. Right, histogram of all extrema detected in the entire MP trace, showing a prominent cluster of peaks around 3800 μV (shaded) and no prominent cluster of negative extrema. As in a, the y-axis is limited. **c**, Left, the first temporal derivative of the MP trace in b shows peaks clearly separated from the noise of the rest of the trace (right). **d**, Left, the second temporal derivative of the MP trace in b also shows clearly separated peaks (right). **e**, For all putative DAP, identified by peaks in the first temporal derivative, the 1^st^ derivative peak and the immediately preceding the 2^nd^ derivative peak are plotted against each other. Data points that extend beyond the noise cloud (black) are identified as DAP (red).

**Fig. S2.**
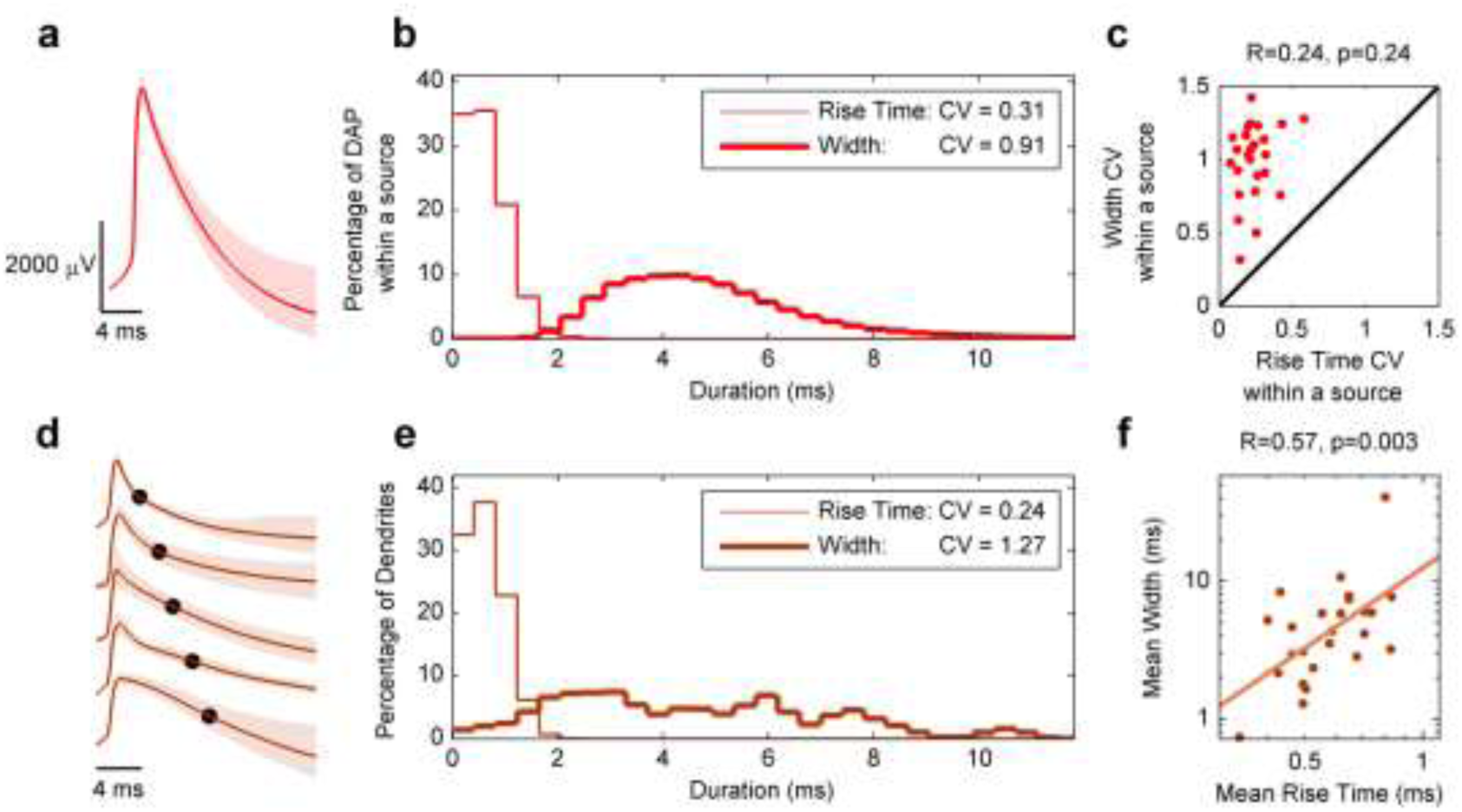
Variability of DAP Width and Rise Time. **a**, Average waveform (median and 25^th^, 75^th^ quantile, n=8917 DAP) for one DAP recording source shows less variation on the rising phase compared to the falling phase. **b**, For a single dendrite source, the rise time (0.49, [0.430, 0.58] ms (median, [25% 75%]), n=67714 DAP, CV=0.31) was smaller (p=0, Wilcoxon signed-rank test) and less variable than the half-width (4.70, [3.59, 6.17] ms (median, [25% 75%]), n=67714 DAP, CV = 0.91). **c**, Across the population of DAP, width CV (1.03, [0.91, 1.15], n=25 dendrites) was always greater (p=1.2×10^−5^, Wilcoxon signed-rank test) than the CV of rise time (0.21, [0.18, 0.26], n=25 dendrites). Width CV and rise time CV were not significantly correlated (r = 0.24, [−0.17, 0.58]; p=0.24, two-sided *t* test). **d**, Average waveforms for five different DAP show a large range of half-widths, marked by black dots. **e**, Across the population of DAP, rise time (0.58, [0.49, 0.65] ms, n=25 dendrites) was significantly shorter (p=2.1×10^−9^, Wilcoxon signed-rank test) than half width (4.33, [2.92, 5.90] ms, n=25 dendrites), and half-width (CV=1.27) was more variable than rise time (CV=0.24). **f**, Across the population of DAP, rise time was significantly correlated with half-width (r=0.57, [0.23, 0.79]; p=2.7×10^−3^, two-sided *t* test). Values in panel b are reported as median and quartile ranges; values elsewhere are reported as median and 95% confidence interval of the median.

**Fig. S3.**
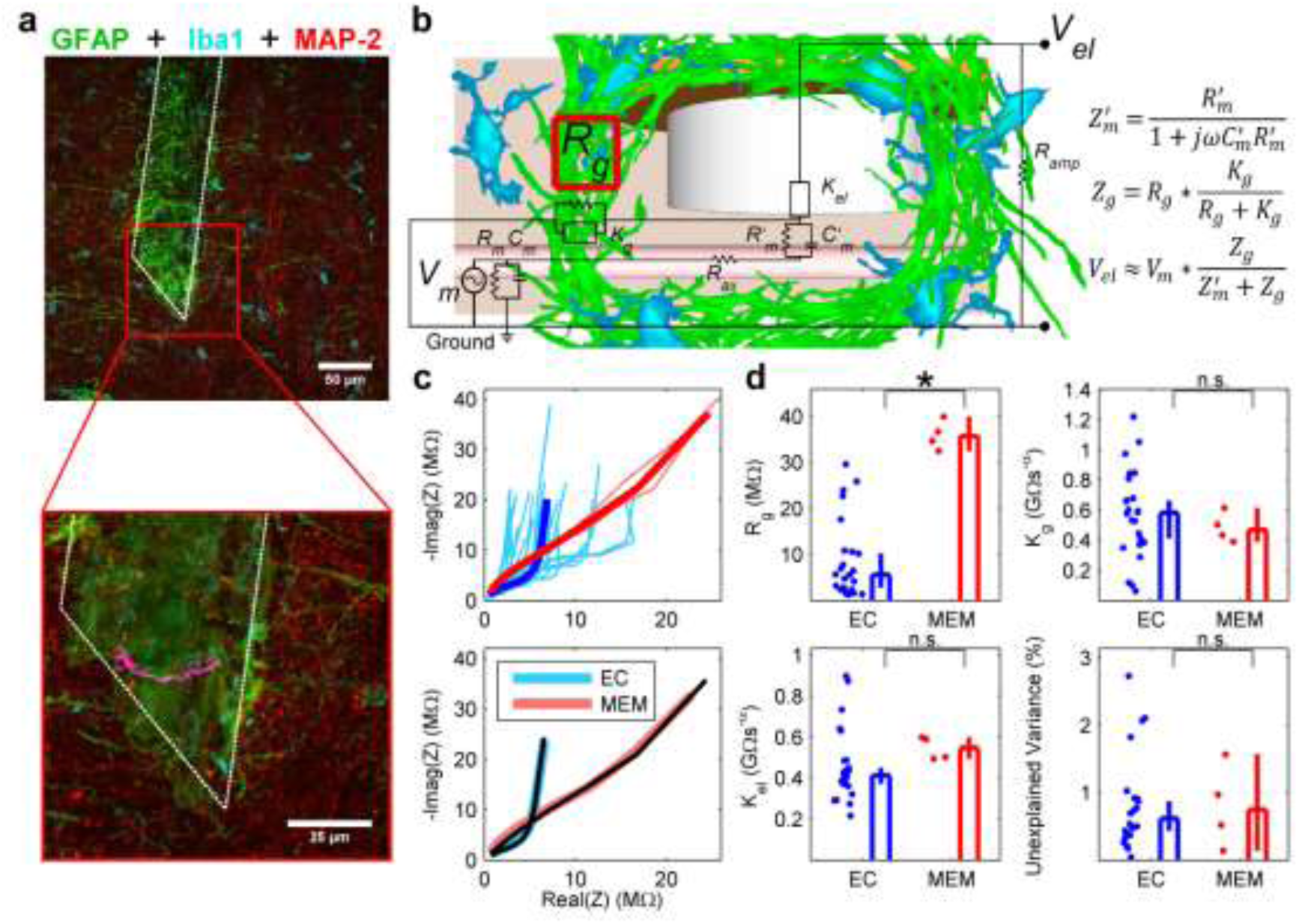
Glial sheath mechanism of DAP recordings. **a**, Top, maximum intensity projection of cortical slice immunohistologically labeled for GFAP (green), Iba1 (cyan), and MAP-2 (red) (see Methods). The white dotted line outlines the putative region once occupied by the tetrode, ~650 μm from the pial surface of Posterior Parietal Cortex. Note the aggregation of GFAP-positive reactive astrocytes encapsulating the tetrode site. Bottom, semi-transparent projection of tetrode tip region. An oblique dendrite segment crossing through the encapsulation area is highlighted in magenta. This dendrite was not recorded, but its proximity to the encapsulated region illustrates the feasibility of the proposed recording mechanism. See Movie S2 for 3-dimensional semi-transparent projection. **b**, Electrical circuit equivalent. The voltage difference between the electrode tip and ground is proportional to the voltage difference between the inside of the dendrite and ground; the magnitude depends on the relative values of the glial sheath impedance *Z_g_* and membrane impedance *Z’_m_*. In typical extracellular recordings, *Z_g_* is negligible compared to *Z’_m_*, so no intracellular signal is recorded. **c**, Top, impedance spectra for normal extracellular (EC, light blue) and DAP-recording (MEM, light red) electrodes shows increased impedance for the DAP-recording electrodes. Dark, bold lines represent the mean impedance spectra. Bottom, fitting the electric circuit model in (b) yields a close approximation (solid black lines) to the mean sample impedance for both extracellular (light blue line) and DAP-recording electrodes (light red line). **d**, The fitted model parameter *R_g_* (glial sheath resistance) for MEM (top left, 35.7, [32.5, 40.0] MΩ) was significantly larger (p=1.8×10^−3^, Wilcoxon rank-sum test) than *R_g_* for EC (5.61, [2.80, 10.2] MΩ), consistent with our proposed glial sheath recording mechanism. In contrast, the fitted model parameters for *K_g_* (top right, EC, 0.58, [0.41, 0.67] GΩ*s^−α^; MEM, 0.47, [0.39, 0.62] GΩ*s^−α^;) and *K_el_* (bottom left, EC, 0.41, [0.37, 0.46] GΩ*s^−α^;; MEM, 0.55, [0.50, 0.60] GΩ*s^−α^;) were not significantly different from each other (*K_g_*, p=0.58; *K_e_i*, p=7.1×10^−2^, Wilcoxon rank-sum test for both).The percentage variance unexplained values (bottom right) for both EC (0.62, [0.43, 0.87] %, n=24 electrodes) and MEM (0.74, [0.14, 1.57] %, n=4 electrodes) were very low, and not significantly different from each other (p=0.87, Wilcoxon rank-sum test).

**Fig. S4.**
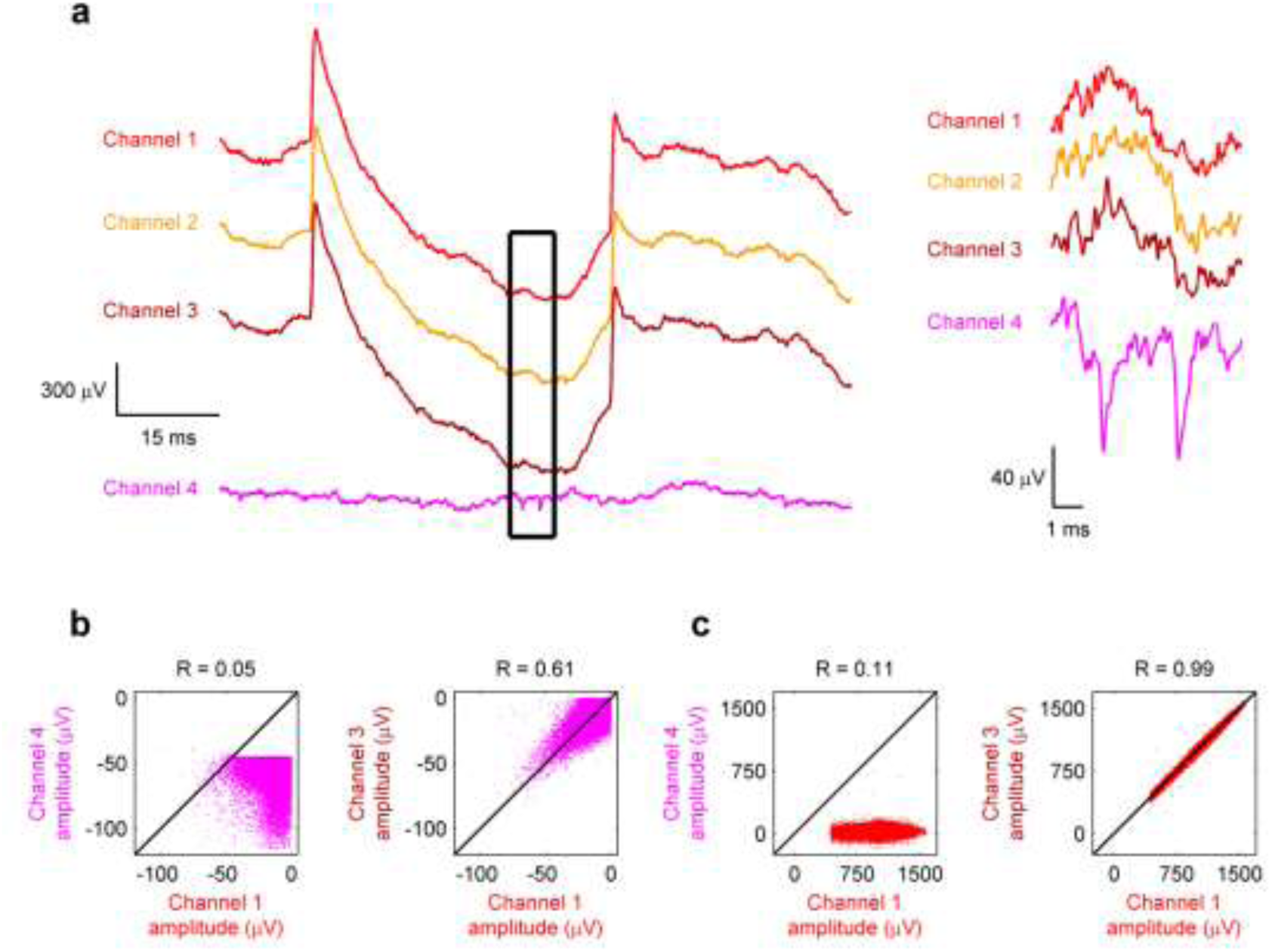
Reduction of MUA on tetrode channels with DMP signal. **a**, Sample trace from a tetrode in which 3 channels recorded DMP (red, orange, brown) but one channel (magenta) did not. No signature of DAP, which are clearly visible on channels 1–3, are visible on channel 4, and no signature of extracellular spikes, which are clearly visible on channel 4 (inset), are visible on channels 1–3. **b**, Extracellular spikes were detected and triggered on channel 4, and the corresponding amplitude recorded on all 4 channels, 2 (out of six possible) combinations of which are shown in scatterplots. Left, channel 4 typically records higher amplitude (more negative) extracellular spikes that are uncorrelated with the amplitude on channels 1–3 (only channels 1 and 4 are shown). Right, extracellular spike amplitude is weakly correlated between channels 1–3 (only channels 1 and 3 are shown). **c**, DAP were detected on channels 1–3, and their amplitude recorded on all 4 channels, 2 combinations of which are shown in scatterplots. Left, channel 4 typically records much smaller amplitude DAP compared to channels 1–3 (only channels 1 and 4 are shown). Right, DAP amplitude on channels 1–3 are highly correlated (only channels 1 and 3 are shown). This shows that the tetrode channel that did not have DAP signal had greater amount of MUA activity, further supporting the glial sheath hypothesis of DAP measurement.

**Fig. S5.**
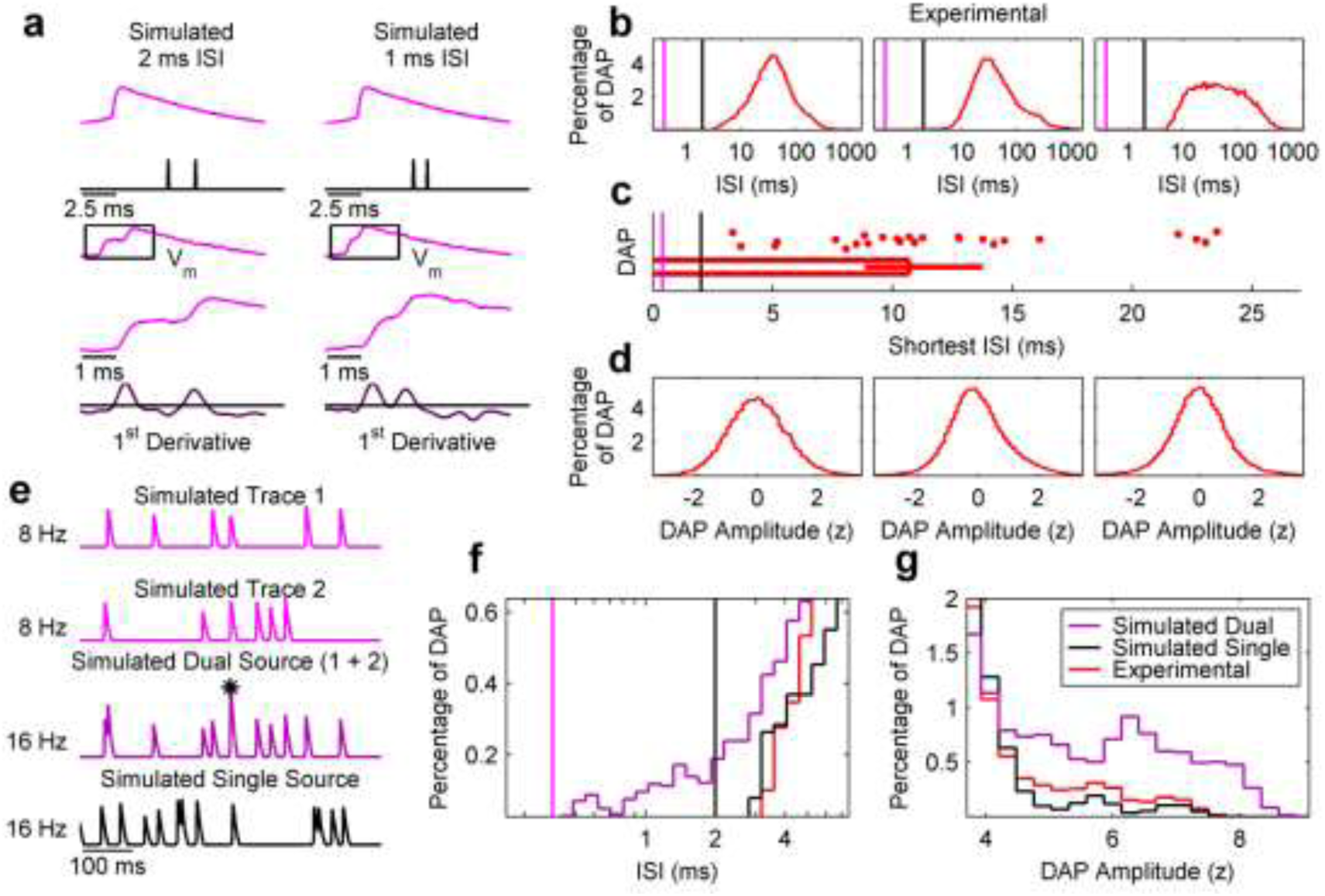
Validation of DAP detection algorithm and separated nature of measured DAP. **a**, Convolution of a typical DAP waveform (top row) with impulses spaced 2 (left), and 1 (right) ms apart. In each row, the y-axis has arbitrary units but identical for the left and right plots. First temporal derivatives of the above traces are plotted in dark purple. In both examples, two peaks above the noise level (see Methods) are clearly distinguishable. **b**, Inter-spike-interval (ISI) histograms of 3 separate experimentally recorded DAP. The black and magenta lines mark the 2 ms interval designating a typical neuronal refractory period and the shortest ISI (0.4 ms) our detection method can distinguish, respectively. **c**, Across the population of DAP, the minimum ISI (10.7, [8.82, 13.8] ms, n=25 dendrites) was significantly longer than the 2 ms refractory period (p=1.2×10^−5^, Wilcoxon signed-rank test), as well as the 0.4 ms resolution limit of our detection method. Data are reported and presented as median and 95% confidence interval of the median. **d**, Unimodal distributions of DAP amplitude are further evidence that each DAP is from a single source. **e**, Two surrogate 8 Hz DAP traces were generated from a gamma process (see Methods) (top two rows, magenta). These were summed together to generate a “Dual Source” 16 Hz DAP trace (third row, purple). The black asterisk indicates an inter-spike interval less than 2 ms. An additional “Single Source” 16 Hz DAP trace was generated from a single 16 Hz gamma distribution (bottom row, black). **f**, Zoomed-in region of the ISI histogram for the simulated dual source16 Hz trace (purple), the simulated single source 16 Hz trace (black), and data from the experimentally recorded DAP represented in the first panel of b (red). The dual source trace has several ISI <2 ms but these are absent in both the single source trace and the experimental DAP trace. **g**, Zoomed-in region of the amplitude histogram for the simulated dual source 16 Hz trace, the simulated single source trace, and the experimental DAP from the first panel of d. The dual source trace has several detected amplitudes greater than 5 standard deviations from the mean of the distribution, but these are absent in both the single source trace and the experimentally observed DAP. These large amplitude events come from the summation of two spikes from two independent simulated 8 Hz sources that occur within 0.4 ms and cannot be resolved; these are not found in experimental data.

**Fig. S6.**
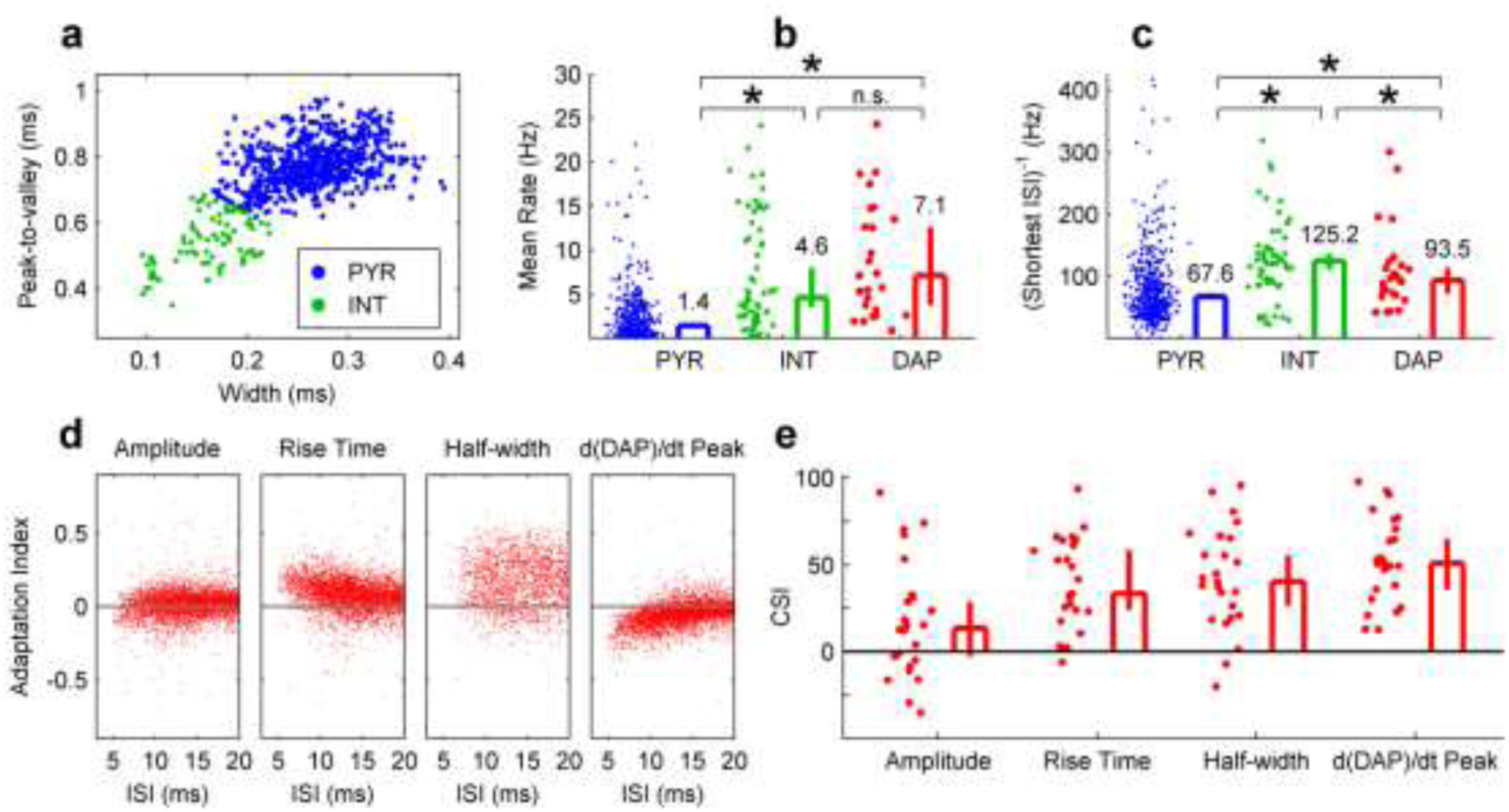
Spiking properties and short term plasticity of pyramidal neurons, interneurons, and DAP in SWS. **a**, Pyramidal neurons and interneurons were identified based on their half-width and the time from the peak of the action potential to the trough after the peak (Peak-to-valley). 87% of extracellularly recorded neurons were classified as pyramidal neurons (657 units), and 13% were classified as interneurons (97 units). **b**, The mean firing rate of pyramidal neurons (1.41, [1.08, 1.65] Hz, n=657 units) was significantly smaller than both that of interneurons (4.59, [3.51, 7.96] Hz, n=97 units; p=7.9×10^−13^, Wilcoxon rank-sum test), and that of DAP (7.07, [3.76, 12.6] Hz, n=25 dendrites; p=1.3×10^−9^, Wilcoxon rank-sum test). DAP mean firing rate was not significantly different from that of interneurons (p=0.31, Wilcoxon rank-sum test). **c**, The peak firing rate of pyramidal neurons (defined as the inverse of lowest 5% of all ISI) (67.6, [63.5, 73.7] Hz, n=657 units) was significantly less than that of both interneurons (125, [111, 137] Hz, n=97 units; p=5.2×10^−9^, Wilcoxon rank-sum test) and DAP (93.5, [72.7, 113] Hz, n=25 dendrites; p=1.0×10^−2^, Wilcoxon rank-sum test). Interneurons also had significantly higher peak firing rates than DAP (p=4.5×10^−2^, Wilcoxon rank-sum test). **d**, Example scatterplots of ISI versus amplitude, rise time, width, and 1^st^ derivative peak demonstrate activity-dependent adaptation. **e**, For the population of DAP, CSI was significantly greater than 0 for measures of amplitude (13.5, [−2.67, 28.8], n=25 dendrites; p=2.6x10^−2^, Wilcoxon signed-rank test), rise time (33.5, [24.2, 58.2], n=25 dendrites; p=2.0×10^−5^, Wilcoxon signed-rank test), half-width (40.3, [26.4, 55.7], n=25 dendrites; p=3.2×10^−5^, Wilcoxon signed-rank test) and 1^st^ derivative peak (50.8, [36.0, 65.0], n=25 dendrites; p=1.4×10^−5^, Wilcoxon signed-rank test)Data are reported and presented as median and 95% confidence interval of the median. * indicates significance at the p<0.05 level, and n.s. indicates lack of significance at the p<0.05 level.

**Fig. S7.**
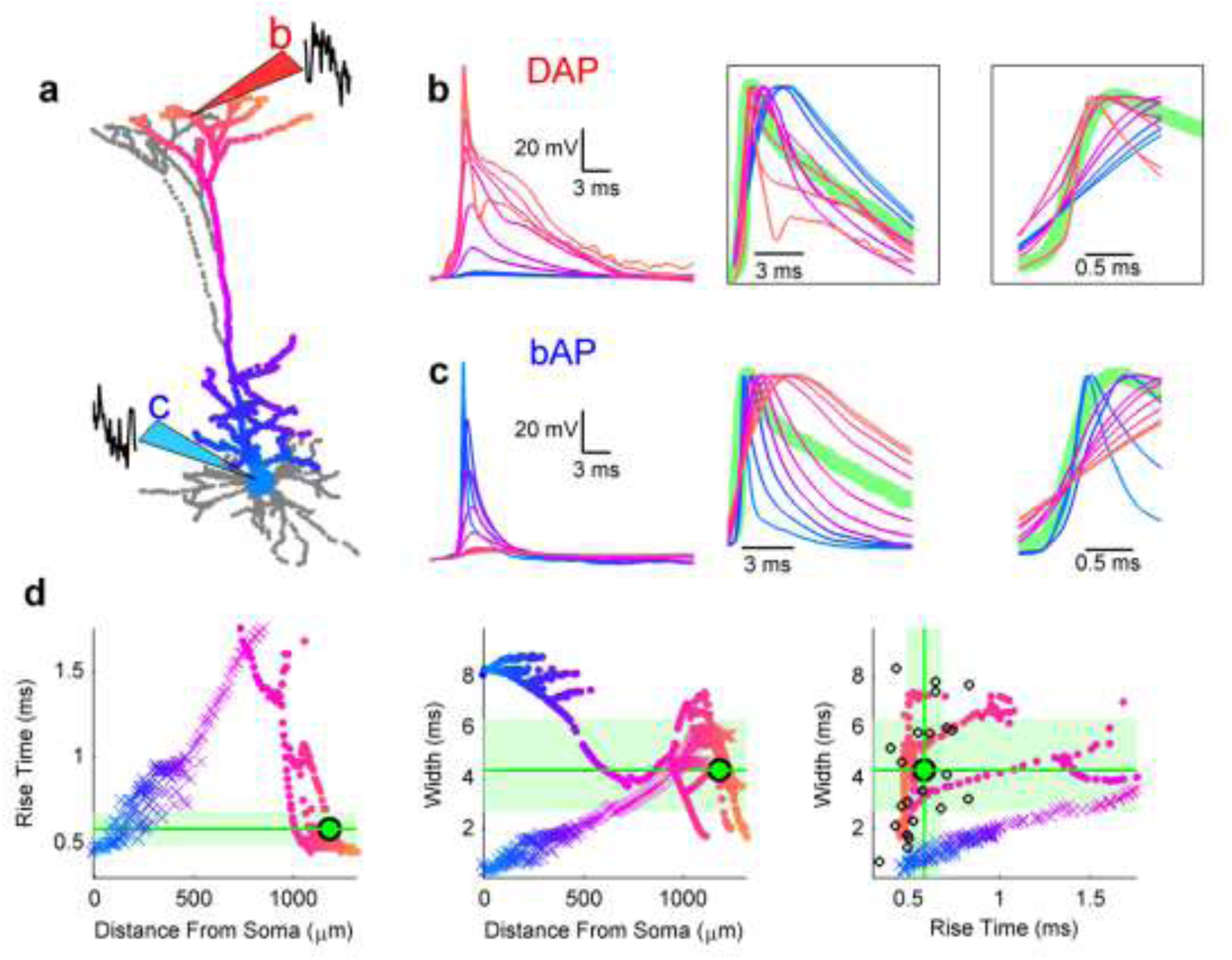
Comparison between experimental measurements of DAP and a biophysical model of layer 5 pyramidal neuron. **a**, A biophysical, multicompartment model of layer 5 pyramidal neuron (see Methods) with small changes in model parameters to fit the in vivo data. The neuron was either stimulated at the soma, or at the distal-most tuft. The apical soma-dendrite axis was mapped on a blue-red color scale for depicting the subsequent figures. **b**, These panels show the results of measuring DAP, initiated at the distal most tufts and then propagating towards the soma. Left panel shows DAP waveform as it travels from the initiation site (large amplitude, red trace) to the soma (very small amplitude blue trace). The DAP amplitude in dendrites is comparable to the estimates from measured data but the somatic signal from DAP is much smaller. To allow the comparison of these waveform shapes across all places on the soma-dendrite axis, the amplitude of each waveform was set to unity (middle). The DAP width increases as it propagates to the soma. The experimentally measured signal’s (green trace) width has a better match with the DAP measured within the distal-most dendrites than with measurements in the soma. To compare the rise time of these results the normalized spike amplitude data is zoomed in (right). This shows that the rise time of experimental data has a better match with DAP in the distal dendrites than elsewhere. **c**, Similar format as in b but for the results of generating a bAP in the soma and measuring its propagation in the dendrites. The bAP amplitude at soma is comparable to the estimates of our experimental measurements (left panel). However, the bAP shape does not match the experimental shape in terms of both width (middle panel) and rise time (right panel) at any place along the soma-dendritic axis. **d**, Left panel: Comparison of bAP (crosses) and DAP (filled circles) rise times at all places along the soma-dendrite axis with the range of values measured experimentally (light green shaded area). The rise time of the example trace in panels b and c is shown by a thick green line and a green dot. bAP measured at a narrow range of proximal dendrites, but DAP measured at a wide range of distal dendrites match the experimental range. Middle, same as in the left panel but for the widths of bAP and DAP versus experimental data. bAP measured at intermediate dendrites, but DAP measured a wide range of distal dendrites overlap with the experimental data. Right, bAP and DAP width time plotted as a function of their rise times. The vast majority simulated DAP rise times and widths, both measured in the distal-most dendrites, match the majority of experimental data. Only a handful of bAP overlap with the experimental data range.

**Fig. S8.**
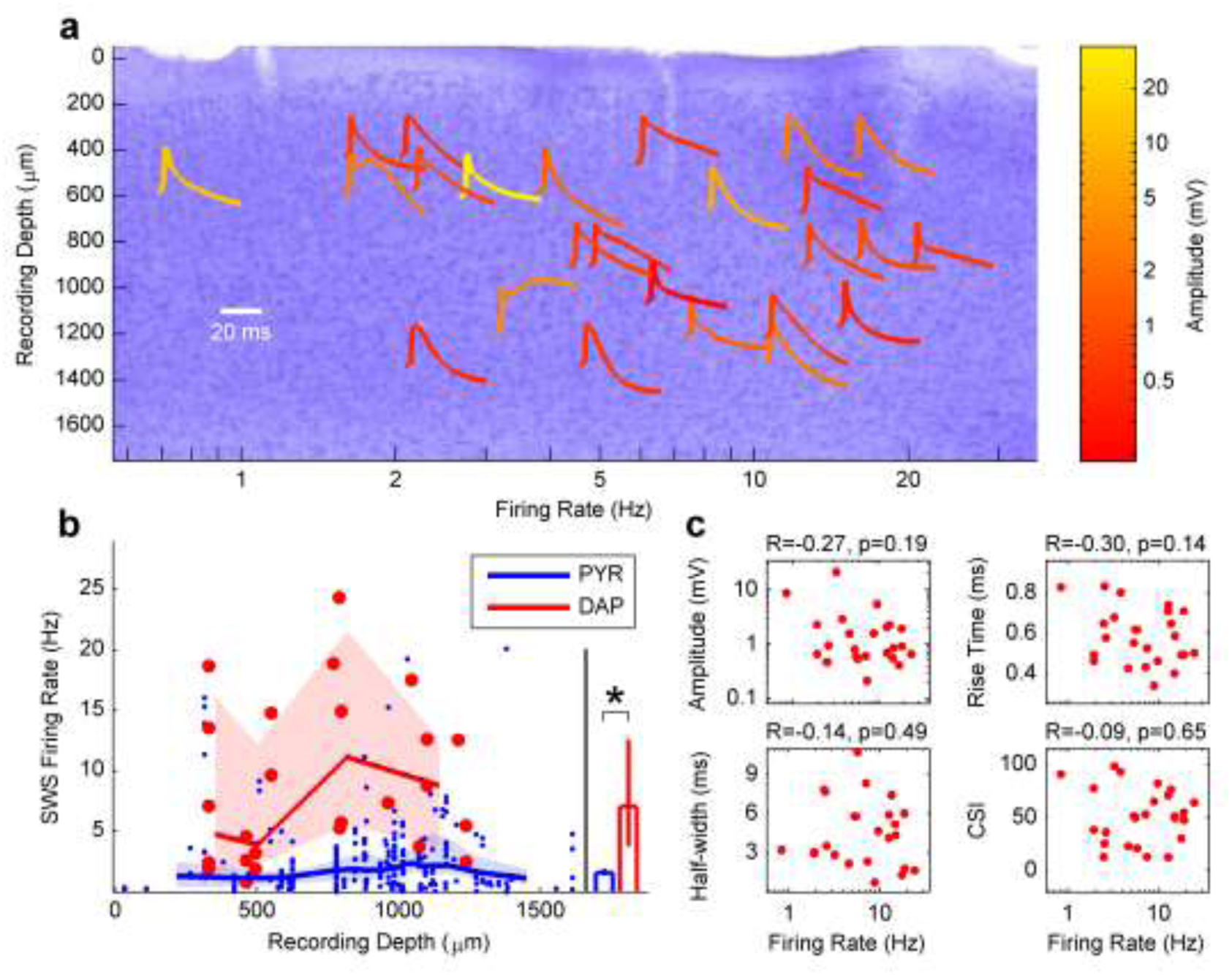
Depth independence of DAP waveform and rate. **a**, Background shows Nissl stained section of a representative parietal cortical tissue from which DAP were recorded. Tetrode tracks are seen are vertical white streaks. Different DAP waveforms, averaged across the respective session, recorded from all animals, are shown as a function of the recording depth and DAP rate. Scale bar shows the duration of the waveforms. Notice that only a handful of DAP waveforms are wide, indicative of calcium spike following the DAP. The majority of DAP are narrow, indicative of only sodium spiking. The DAP rate was not significantly correlated with recording depth (770, [460, 960] μm, n=25 DAP sources; r=0.24, [−0.17, 0.58]; p=0.24, two-sided *t* test). **b**, The average DAP rate was computed as a function of the recording depth across all data. This was compared with the average firing rates of ensembles of pyramidal neurons' somatic spikes as a function of depth, measured under similar conditions as DAP. DAP rates were far greater than pyramidal neuron somatic spike rates at all depths (p=3.7×10^−4^ for group effect, two-way ANOVA with recording depth as a continuous predictor). **c**, DAP mean firing rate was not significantly correlated with amplitude (r=−0.31, [−0.63, 0.10]; p=0.13, two-sided *t* test), rise time (r=0.22, [−0.19, 0.56]; p=0.30, two-sided *t* test), half-width (r=−0.07, [−0.45, 0.34]; p=0.76, two-sided *t* test), or CSI (r=−0.31, [−0.63, 0.10]; p=0.14, two-sided *t* test).

**Fig. S9.**
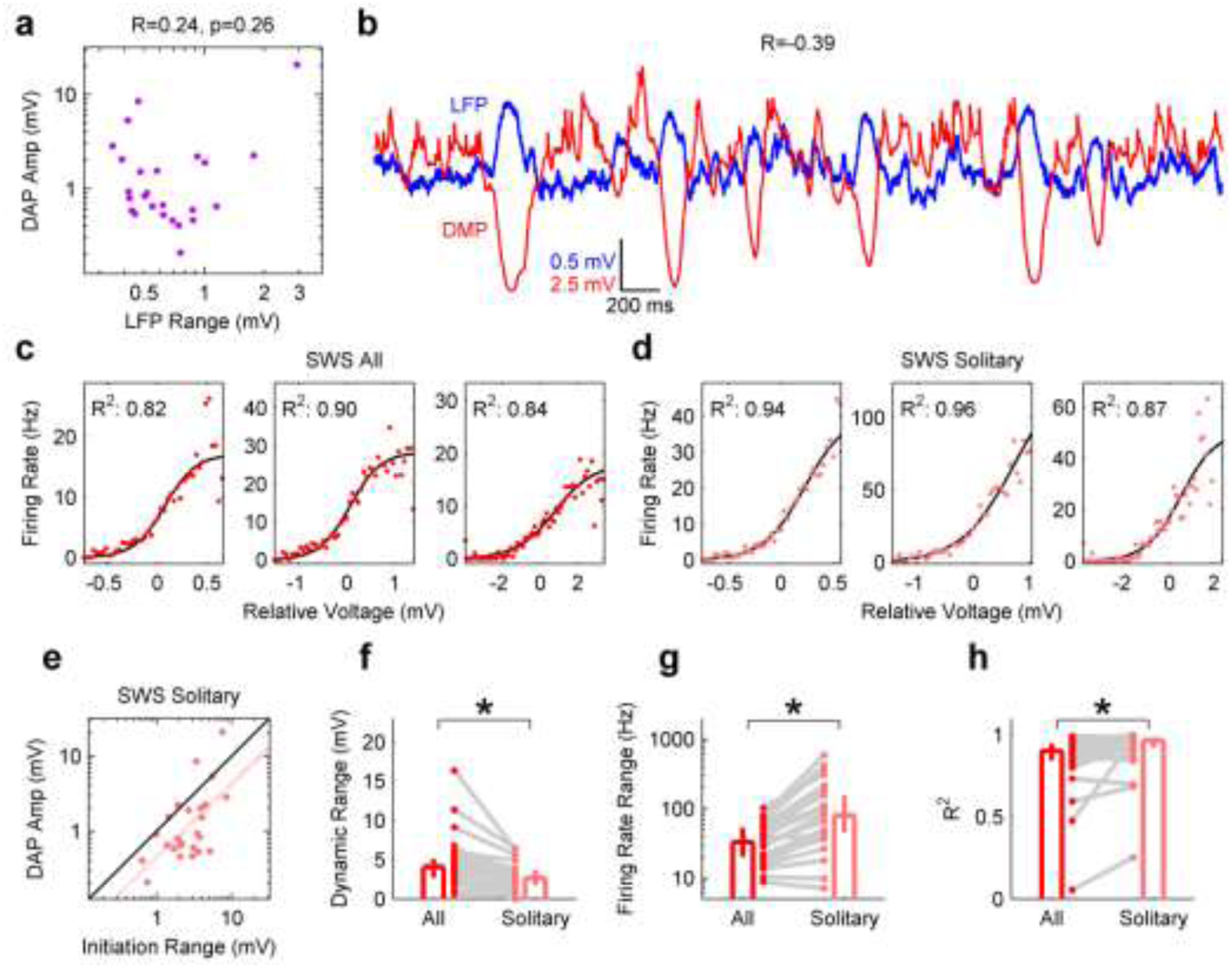
Modulation of DAP by subthreshold membrane potentials. **a**, DAP amplitude was not significantly correlated (r=0.24, [−0.18, 0.58]; p=0.26, two-sided *t* test) with the range of LFP recorded simultaneously from a nearby tetrode, ruling out spurious noise artifacts. **b**, During SWS, simultaneously recorded LFP (blue) and DMP (red) show up-down states with reversed polarity with respect to each other. **c**, Sample Voltage-Rate (V-R) curves computed using all DAP within a session during SWS. **d**, Sample V-R curves for the same DAP in C, but for only those DAP and times separated from other DAP by at least 50 ms (Solitary). **e**, For solitary DAP in SWS, initiation range (2.99, [1.94, 3.73] mV, n=25 dendrites) was larger (p=3.6×10^−4^, Wilcoxon signed-rank test) than the corresponding DAP amplitude (0.83, [0.60, 1.90] mV, n=25 dendrites), and positively correlated (r=0.61, [0.28, 0.81], p=1.2×10^−3^, two-sided *t* test). **f**, The dynamic voltage range for solitary DAP (2.66 [1.75, 3.71], n=25 dendrites) was smaller (p=1.7×10^−2^, Wilcoxon signed-rank test) than for all DAP (4.03, [2.74, 5.07], n=25 dendrites). **g**, The V-R firing rate range was significantly higher (p=5.9×10^−3^, Wilcoxon signed-rank test) for solitary DAP (79.8, [45.5, 154] Hz, n=25 dendrites) compared to all DAP (33.1, [20.3, 53.1] Hz, n=25 dendrites). h, The goodness of the logistic fit of the VR-curve when calculated for solitary DAP (0.97, [0.92, 0.97], n=25 dendrites), was slightly larger (p=2.0×10^−2^, Wilcoxon signed-rank test) compared to all DAP (0.90, [0.84, 0.95], n=25 dendrites). Data are reported and presented as median and 95% confidence interval of the median, and * indicates significance at the p<0.05 level.

**Fig. S10.**
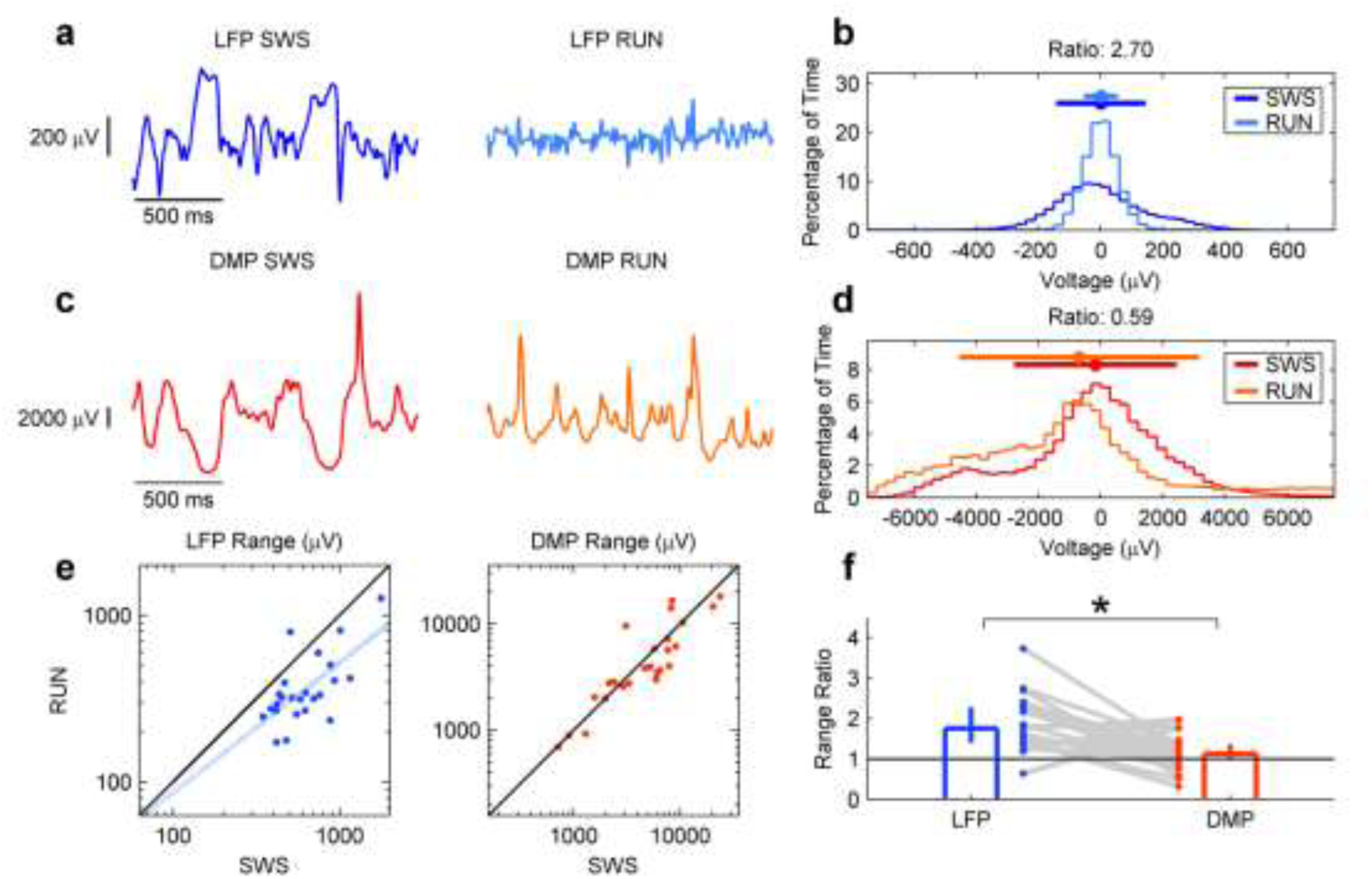
Comparison of LFP and subthreshold DMP modulation during SWS and RUN. **a**, Sample segments of a spike-clipped LFP trace during SWS (left, blue) and RUN (right, light blue), showing smaller amplitude fluctuations during RUN. **b**, Histograms of the LFP voltage for the traces in a, showing a much wider range of variation in SWS (−265 to 215 μV, range of 480 μV) than during RUN (−89.7 to 87.9 μV, range of 178 μV). The ratio of SWS range to RUN range in this example is 2.70, indicating a much larger range in SWS. **c**, Sample segments of a spike-clipped MP trace during SWS (left, red) and RUN (right, orange), showing large amplitude fluctuations in both SWS and RUN. **d**, Histograms of the MP voltage for the traces in c, showing a similar range of variation in SWS (−4690 to 3560 μV, range of 8250 μV) and RUN (−6080 to 7980 μV, range of 14100 μV). The ratio of SWS range to RUN range in this example is 0.59, indicating a comparable range in SWS and RUN. **e**, Left, spike-clipped local field potential range during RUN (320, [275, 407] μV, n=25 recording segments here and throughout the figure) was smaller (p=4.6×10^−5^, Wilcoxon signed-rank test) than the local field potential range during SWS (571, [468, 761] μV); the two ranges were also significantly correlated (r=0.74, [0.49, 0.88], p=2.4×10^−5^, two-sided *t* test). Right, spike-clipped DMP range during RUN (3820, [2760, 6170] μV) was not significantly different (p=9.3×10^−2^, Wilcoxon signed-rank test) than that during SWS (5720, [2920, 7690] μV), and the two measures were significantly correlated (r=0.89, [0.76, 0.95], p=3.7×10^−9^, two-sided *t* test). **f**, The ratio of SWS range to RUN range in the LFP (1.75, [1.41, 2.28], n=25 recording segments) was significantly greater than 1 (p=2.3×10^−5^, Wilcoxon signed-rank test) and significantly greater (p=5.9×10^−5^, Wilcoxon signed-rank test) than that of DAP (1.13, [1.00, 1.36]), which was not significantly different from 1 (p=6.1×10^−2^, Wilcoxon signed-rank test).

**Fig. S11.**
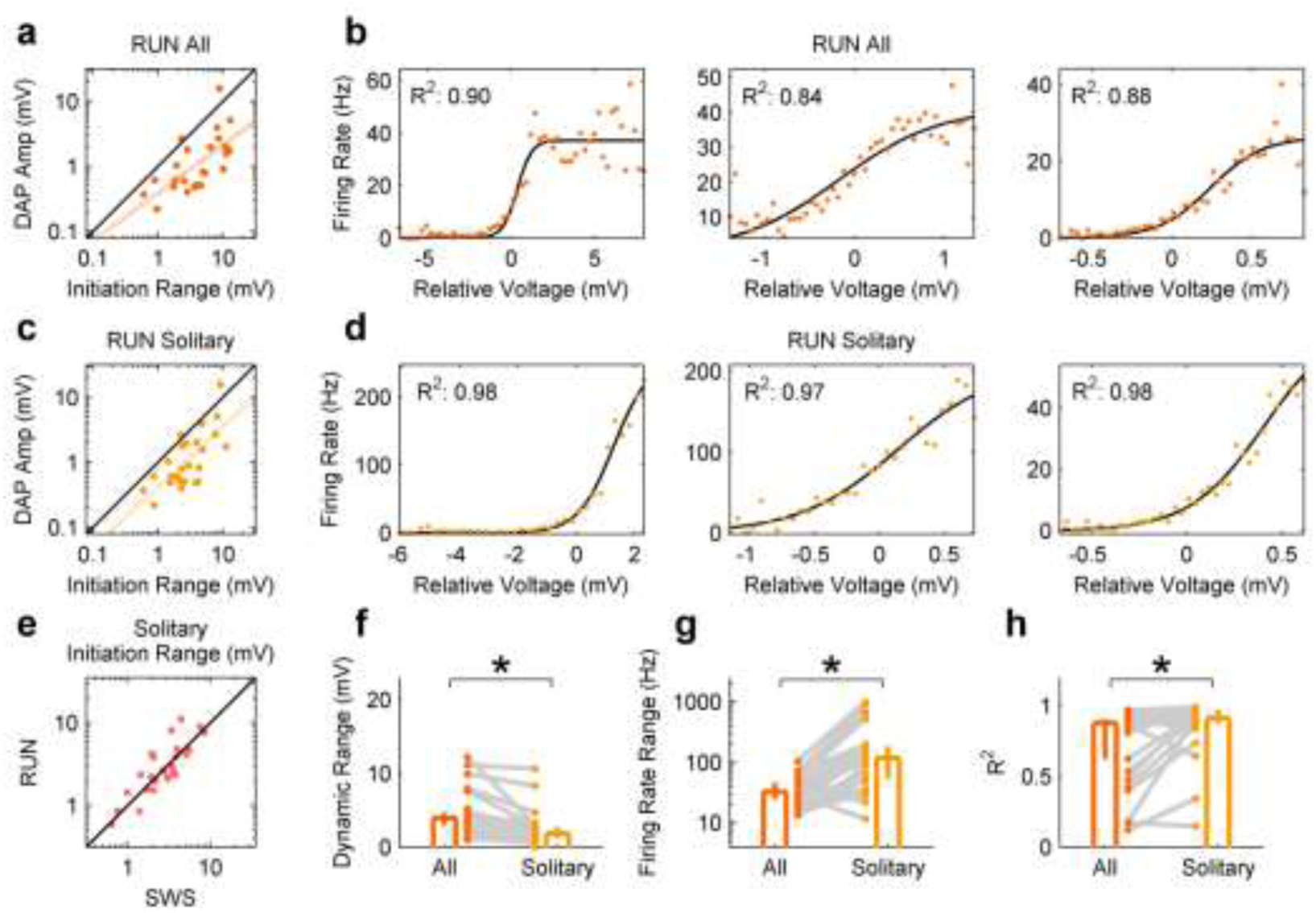
Properties of solitary DAP in SWS and RUN. **a**, DAP initiation range in RUN (4.07, [2.51, 8.14] mV, n=25 dendrites) was larger (p=2.5x10^−5^, Wilcoxon signed-rank test) than the corresponding DAP amplitude (0.82, [0.52, 1.78] mV, n=25 dendrites), and positively correlated (r=0.65, [0.34, 0.83], n=25 dendrites; p=4.3×10^−4^, two-sided *t* test) **b**, Sample Voltage-Rate (V-R) curves for all DAP during RUN. **c**, For solitary DAP in RUN, initiation range (2.56, [2.14, 4.13] mV, n=25 dendrites) was larger (p=9.4×10^−5^, Wilcoxon signed-rank test) than the corresponding DAP amplitude, and positively correlated (r=0.70, [0.42, 0.86]; p=9.4×10^−5^, two-sided *t* test). **d**, Sample V-R curves for the same DAP in (b), but for only those DAP and times separated from other DAP by at least 50 ms. **e**, For solitary DAP, initiation range in SWS (2.99, [1.94, 3.73] mV, n=25 dendrites) and RUN (2.56, [2.14, 4.13] mV, n=25 dendrites) were positively correlated (r=0.84, [0.66, 0.93]; p=1.7×10^−7^, two-sided *t* test) and not significantly different (p=0.74, Wilcoxon signed-rank test). **f**, As in Figure 5g, 24 of 25 dendrites had sufficient data to characterize V-R curves in RUN. The dynamic range for solitary DAP in RUN (1.75, [1.25, 2.69] mV, n=24 dendrites) was slightly reduced (p=1.8×10^−3^, Wilcoxon signed-rank test), compared to all DAP in RUN (3.90, [2.98, 4.67] mV, n=24 dendrites). **g**, The V-R firing rate range was significantly higher (p=2.3×10^−4^, Wilcoxon signed-rank test) for solitary DAP in RUN (117, [52.6, 183] Hz, n=24 dendrites) compared to all DAP in RUN (32.5, [24.6, 46.5] Hz, n=24 dendrites). **h**, The V-R curve in RUN was better approximated by the logistic fit (p=3.8×10^−2^, Wilcoxon signed-rank test) when calculated for solitary DAP (0.91, [0.88, 0.97], n=24 dendrites) compared to all DAP (0.88, [0.62, 0.90], n=24 dendrites).

**Fig. S12.**
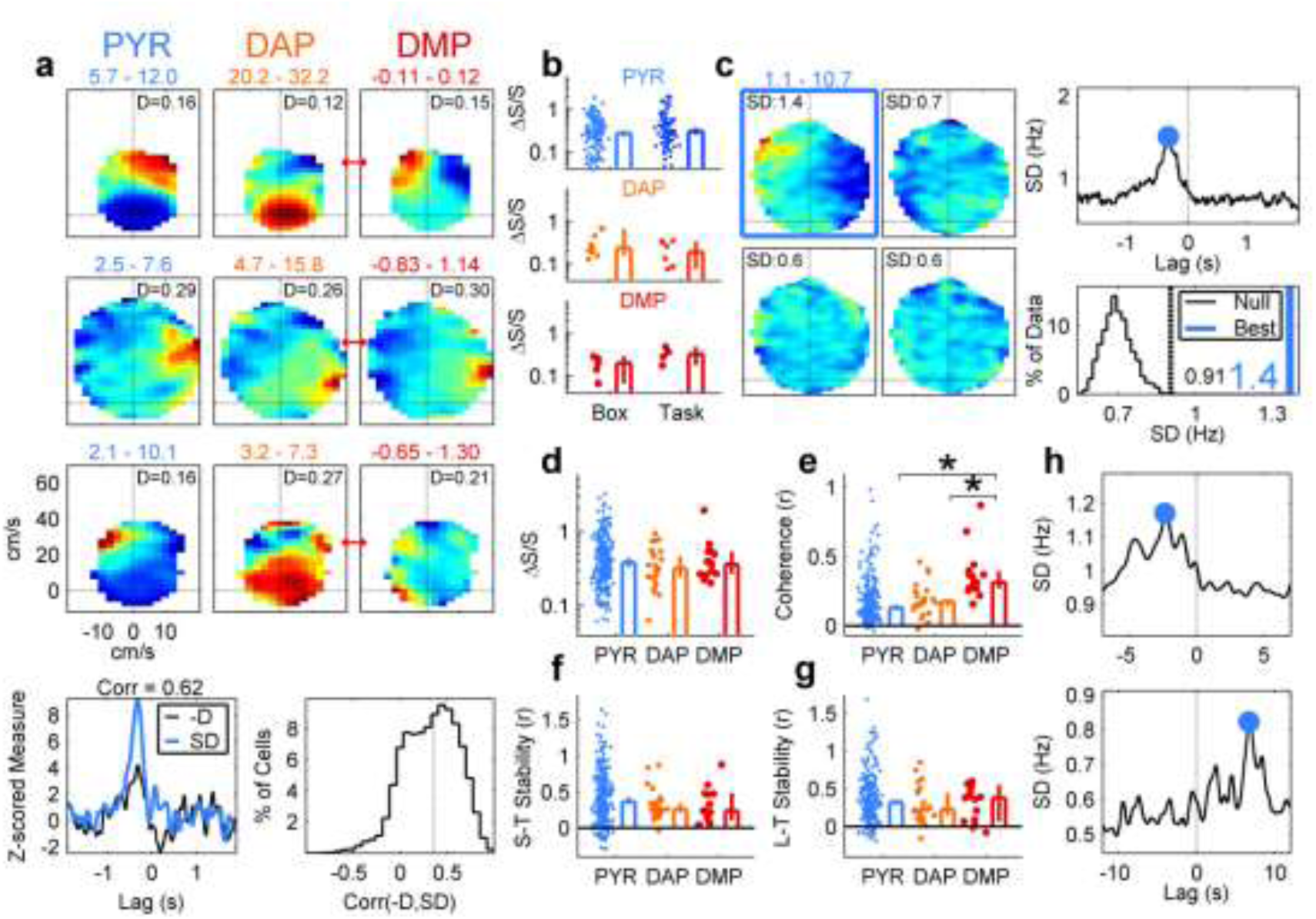
Sample egocentric rate maps, shuffle procedure, additional measures, long-term stability. **a**, Top three rows, three sample pyramidal soma (PYR, left), DAP (middle), and DMP (right) egocentric maps. Bottom row, dispersion (D) and standard deviation (SD) are highly correlated as a function of lag for a sample unit (left) and for the entire population. The higher signal-to-noise ratio of SD motivates its use in further analyses. **b**, There were no significant differences in normalized standard deviation (see panels c, d, and Methods) for units recorded in the sleep box or during the random foraging task, for PYR, DAP, or DMP. **c**, Illustration of shuffling method used to determine significance and optimal lag (see Methods). The standard deviation (SD) of the time-shifted maps (top) is plotted as a function of lag (bottom-left). The peak SD (blue mark) is compared to the distribution of SDs at long time-lags (black, bottom-right). **d**, Normalized standard deviation ΔS/S, see Methods), was comparable between PYR (0.38, [0.33, 0.44], n=245 maps), DAP (0.32, [0.25, 0.46], n=24 maps), and DMP (0.36, [0.27, 0.53], n=15 maps), with no significant differences (PYR vs DAP, p=0.31; PYR vs DMP, p=0.66; DAP vs DMP, p=0.74, Wilcoxon rank-sum test for all). **e**, Coherence for both PYR (0.13, [0.11, 0.15, n=245 maps) and DAP (0.17, [0.14, 0.19], n=24 maps) was significantly smaller than coherence of DMP (0.31, [0.27, 0.39], n=15 maps; PYR vs. DMP, p=3.9×10^−6^; DAP vs DMP, p=3.6×10^−5^, Wilcoxon rank-sum test for both); DAP and PYR map coherence were not different from each other (p=0.33, Wilcoxon rank-sum test). **f**, Short-term (S-T) stability (see Methods) for PYR (0.37, [0.33, 0.43], n=245 maps) was slightly higher than DAP (0.26, [0.21, 0.34], n=24 maps) and DMP (0.24, [0.10, 0.48], n=15 maps), but these differences were not statistically significant (PYR vs DAP, p=0.13; PYR vs DMP, p=0.16, Wilcoxon rank-sum test for both), nor was the difference between DAP and DMP (p=0.61, Wilcoxon rank-sum test). **g**, Long-term (L-T) stability (see Methods) for pyramidal soma (0.33, [0.29, 0.36], n=245 maps) was comparable (p=0.25, Wilcoxon rank-sum test) to that of DAP (0.23, [0.15, 0.45], n=24 maps), and DMP (0.38, [0.07, 0.55], n=15 maps; p=0.76, Wilcoxon rank-sum test). DAP and DMP L-T stability were not significantly different (p=0.92, Wilcoxon rank-sum test). **h**, Two sample pyramidal neurons with an extremely large lag time of maximal standard deviation, at −2.4 s (top) and 6.7 s (bottom) respectively.

**Table S1.**
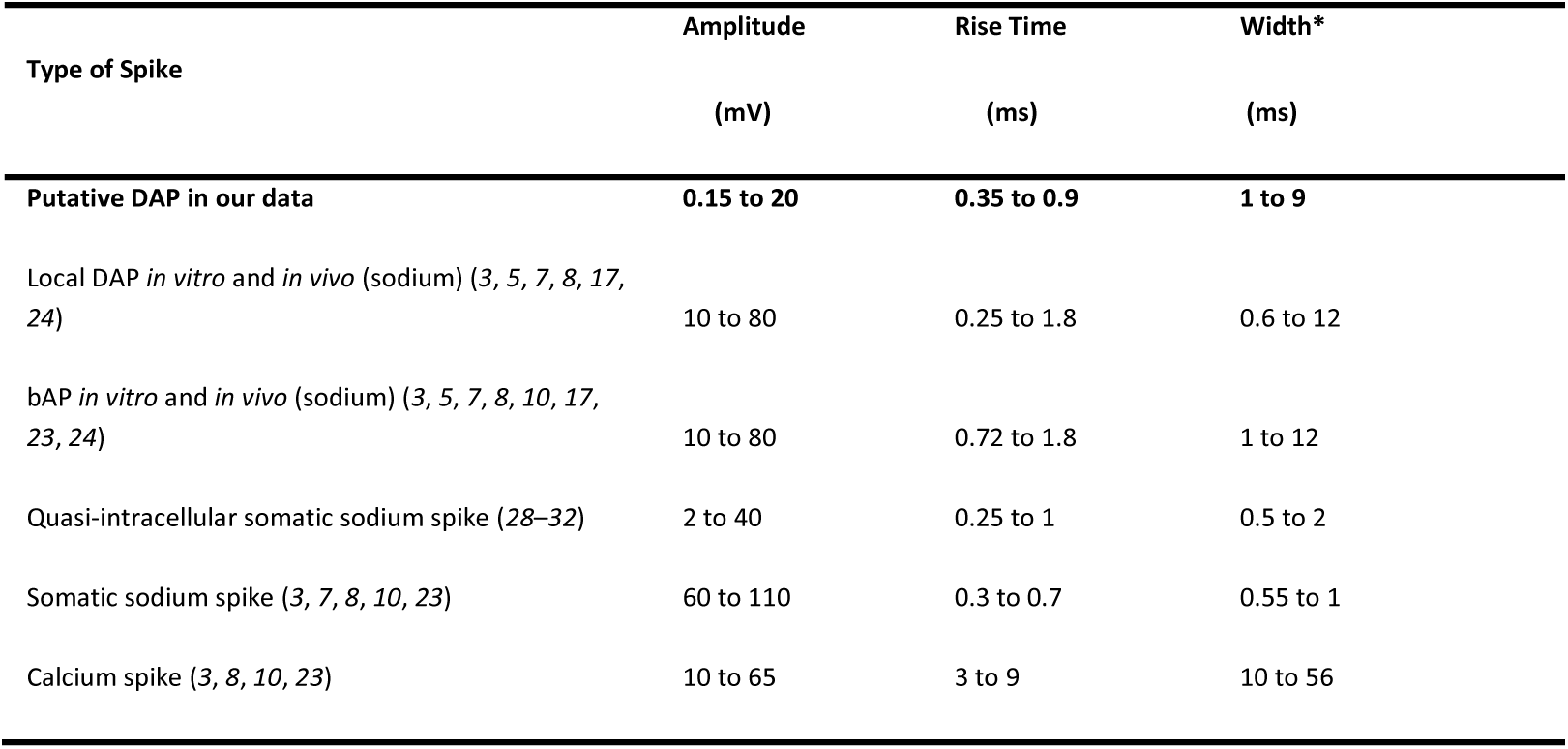
Characterization of the amplitude, rise time, and width for putative DAP from the current study compared to the same measures for different types of intracellularly-recorded spikes. Note that our data have widths approximately an order of magnitude larger than somatic sodium spikes but an order of magnitude smaller than calcium spikes, making these unlikely explanations for our data, but match dendritic sodium spike properties. *: *“Width” refers to width at the base for calcium spikes, and width at half maximum for all others*.

#### Movie S1

**Sample voltage traces of simultaneously-recorded local field potential and dendritic membrane potential**. Voltage traces show dendritic membrane potential (DMP) recorded on a tetrode (bottom, red) simultaneously recorded with the cortical local field potential (LFP) on a nearby tetrode (top, blue). Both tetrodes are in the parietal cortex. The sound accompanying the video is a direct translation of the two waveforms, and is in stereo: the left channel is the sound of the LFP and the right channel is the sound of the DMP. Note the different voltage scales for the two traces, demonstrating the large amplitude and inverted polarity of dendritic action potentials (DAP) compared to extracellular spikes.

#### Movie S2

**3-dimensional semi-transparent projection of immunohistologically stained cortical slice**. Slice (same image stack as in Extended Data Figure 3a, bottom) is labeled for GFAP (green, putative reactive astrocytes), Iba1 (cyan, putative microglia), and MAP-2 (red, putative dendrites). An oblique dendrite segment trapped within the glial sheath is highlighted in magenta. The dense aggregation of GFAP and Iba1 around the tetrode represents the glial sheath formed by astrocytes and microglia in response to the implanted tetrode.

